# CellOracle: Dissecting cell identity via network inference and in silico gene perturbation

**DOI:** 10.1101/2020.02.17.947416

**Authors:** Kenji Kamimoto, Christy M. Hoffmann, Samantha A. Morris

## Abstract

Here, we present CellOracle, a computational tool that integrates single-cell transcriptome and epigenome profiles to infer gene regulatory networks (GRNs), critical regulators of cell identity. Leveraging inferred GRNs, we simulate gene expression changes in response to transcription factor (TF) perturbation, enabling network configurations to be interrogated *in silico*, facilitating their interpretation. We validate the efficacy of CellOracle to recapitulate known regulatory changes across hematopoiesis, correctly predicting the outcomes of well-characterized TF perturbations. Integrating CellOracle analysis with lineage tracing of direct reprogramming reveals distinct network configurations underlying different reprogramming failure modes. Furthermore, analysis of GRN reconfiguration along successful reprogramming trajectories identifies new factors to enhance target cell yield, uncovering a role for the AP-1 subunit Fos, with the hippo signaling effector, Yap1. Together, these results demonstrate the efficacy of CellOracle to infer and interpret cell-type-specific GRN configurations, at high-resolution, promoting new mechanistic insights into the regulation and reprogramming of cell identity.

## Introduction

Defining the transitions between cellular identities and states is central to our understanding of development and disease. Gene Regulatory Networks (GRNs) represent the complex, dynamic molecular interactions that act as critical determinants of cell identity. These networks describe the intricate interplay between transcriptional regulators and multiple cis-regulatory DNA sequences, resulting in the precise spatial and temporal regulation of gene expression (Davidson and Erwin, 2006). GRN hierarchy is established during development, creating a primary spatial organization to support faithful tissue patterning; successive GRN implementation initiates the formation of regional cell identities, leading to steady-state expression programs that anchor terminally differentiated cell identities (Peter and Davidson, 2011, 2016). Systematically delineating GRN structures enables a logic map of regulatory factor cause-effect relationships to be mapped (Materna and Davidson, 2007). In turn, this knowledge supports a better understanding of how cell identity is determined and maintained, informing new strategies for cellular reprogramming to support disease modeling or cell-based therapeutic approaches.

Traditionally, GRN reconstruction has required painstaking experimentation (Materna and Oliveri, 2008). Recently, genomic technologies have enabled the collection of large-scale transcriptome and epigenome data, paving the way for computational prediction of GRNs (Chai et al., 2014; Thompson et al., 2015). For example, our previous CellNet platform used microarray data from bulk populations to reconstruct GRNs, supporting a systematic assessment of cell identity and prioritization of factors to enhance cell reprogramming (Cahan et al., 2014; Morris et al., 2014). However, the use of bulk expression data to infer networks from heterogenous populations obscured crucial differences between cell sub-types and altogether masked rare cell types. Single-cell genomics, now representing a range of modalities to capture transcriptome and epigenome data, enables this population heterogeneity to be deconstructed (Stuart and Satija, 2019). The emergence of single-cell technologies has been accompanied by methods to infer GRNs from the associated high-dimensionality datasets (Fiers et al., 2018).

Nonetheless, inferring GRNs from gene expression data is a challenging task; most GRN inference algorithms use correlation to infer regulatory connections, but correlation does not imply causation. Indeed, evaluation of several computational methods to reconstruct networks from single-cell data has demonstrated their poor performance (Chen and Mar, 2018; Pratapa et al., 2020), likely due to high levels of noise and drop-out (where expressed genes are undetected by single-cell RNA-sequencing, scRNA-seq). Some methods have aimed to tackle these limitations (Aibar et al., 2017; Iacono et al., 2019), but it remains a challenge to infer GRNs from single-cell data. Perhaps even more pertinent is the lack of methodologies to interpret the resulting networks, hindering valuable biological insight into the relationship between GRN and cell phenotype.

Here, we present CellOracle, a machine learning-based tool to infer GRNs via the integration of different single-cell data modalities. CellOracle overcomes current challenges in GRN inference by using single-cell transcriptomic and chromatin accessibility profiles, integrating prior biological knowledge via regulatory sequence analysis to infer transcription factor (TF)-target gene interactions. Moreover, we designed CellOracle to apply inferred GRNs to the simulation of gene expression changes in response to TF perturbation. This unique feature enables inferred GRN configurations to be interrogated *in silico*, facilitating their interpretation. Here, we benchmark CellOracle against ground-truth TF-gene interactions, outperforming existing algorithms, and demonstrate its efficacy to recapitulate known regulatory changes across hematopoiesis, correctly predicting well-characterized phenotypic changes in response to TF perturbations.

Furthermore, we apply CellOracle to interrogate GRN reconfiguration during the direct lineage reprogramming of fibroblasts to induced endoderm progenitors (iEPs), a prototypical TF-mediated fate conversion protocol. These analyses, together with our previous lineage-tracing strategy, reveal distinct modes of reprogramming failure, defined by unique GRN signatures that emerge at the early stages of conversion. Using principles of graph theory to identify critical nodes in conjunction with *in silico* simulation reveals previously undescribed roles for several TFs in reprogramming. We experimentally validate these predictions via TF overexpression and Perturb-seq-based knockout. We also demonstrate that one of these TFs, *Fos*, plays roles in both iEP reprogramming and maintenance, where interrogation of inferred *Fos* targets highlights a putative role for AP1-Yap1 in fibroblast to iEP conversion. Together, these results demonstrate the efficacy of CellOracle to infer and interpret cell-type-specific GRN configurations at high-resolution, enabling new mechanistic insights into the regulation and reprogramming of cell identity. CellOracle code and documentation are available at https://github.com/morris-lab/CellOracle.

## Results

### Construction of CellOracle for GRN Inference

In this study, we aimed to develop a computational approach to identify critical regulators of cell differentiation and reprogramming. To achieve this, we infer GRN configurations to reveal how networks are rewired during the establishment of defined cellular identities and states, highlighting known and putative regulatory factors of fate commitment. CellOracle overcomes population heterogeneity by leveraging single-cell genomic data, enabling accurate inference of the GRN dynamics underlying complex biological processes. This approach offers higher resolution, relative to building a universal GRN for each terminal cell identity of interest. Moreover, we aim to overcome current challenges by dividing GRN inference into stepwise tasks, integrating different data modalities at each stage (**Figure 1A-D; methods**). Central to this approach is our identification and use of accessible promoter/enhancer DNA sequences, in concert with TF binding motifs to define the directionality of gene-gene connections and minimize false-positive edges. This strategy contrasts with the inference of causality from gene expression measurements alone, which can be problematic, particularly for relatively noisy single-cell datasets (Kim et al., 2015; Pratapa et al., 2020) (**Figure 1A**).

**Figure 1.**
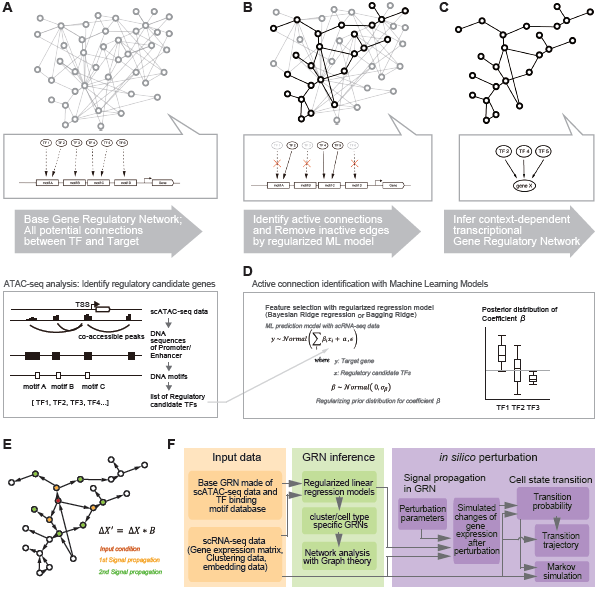
Construction of CellOracle. **(A)** Overview of the CellOracle pipeline to infer context-dependent GRN configurations for transcriptionally-defined cell types. First, genomic DNA sequence and TF binding motif information provide all potential regulatory links to construct a ‘base GRN’ (Upper panel). In this step, CellOracle uses scATAC-seq data to identify accessible promoter/enhancer DNA sequences. The DNA sequence of regulatory elements is scanned for TF binding motifs, generating a list of potential regulatory connections between a TF and its target genes (Lower panel). **(B)** Active connections (described below), which are dependent on cell state or cell type, are identified from all potential connections in the base GRN. **(C)** Cell type- and state-specific GRN configurations are constructed by pruning insignificant or weak connections. **(D)** For steps **C** and **D**, using single-cell expression data, an active connection between the TF and the target gene is identified for defined cell identities and states by building a machine learning (ML) model that predicts the relationship between the TF and the target gene. ML model fitting results present the certainty of connection as a distribution, enabling the identification of GRN configurations by removing inactive connections from the base GRN structure. **(E)** Schematic illustration of the CellOracle approach for simulating future gene expression after perturbation. Leveraging the features of the linear predictive ML model, CellOracle simulates how target gene expression changes in response to the changes in regulatory gene expression. Iterative calculation enables the estimation of indirect downstream effects resulting from the perturbation of a single TF. **(F)** Flowchart of the CellOracle workflow.

In the first step of the CellOracle pipeline, single-cell chromatin accessibility data (scATAC-seq) is used to assemble a ‘base’ GRN structure, representing a list of all potential regulatory genes that are associated with each defined DNA sequence. This step leverages the transcriptional start site (TSS) database (http://homer.ucsd.edu/homer/ngs/annotation.html), and Cicero, an algorithm that identifies co-accessible scATAC-seq peaks (Pliner et al., 2018), to identify accessible promoters/enhancers. The DNA sequence of these regulatory elements is then scanned for TF binding motifs, repeating this task for all regulatory sequences, to generate a base GRN structure of all potential regulatory interactions (**Figure 1A**). Here, we exclusively use scATAC-seq data to assemble base GRN structures. However, any other genomic data that includes regulatory DNA sequence information, such as bulk ATAC-seq, DNase-seq, and Hi-C, can potentially be integrated into the CellOracle pipeline. Here, we demonstrate CellOracle by constructing a base GRN using a published mouse scATAC-seq atlas consisting of ∼100,000 cells across 13 tissues, representing ∼400,000 differentially-accessible elements, and 85 different chromatin patterns (Cusanovich et al., 2018). This base GRN is built into the CellOracle library, to support GRN inference in the absence of sample-specific scATAC-seq datasets.

The second step in the CellOracle pipeline uses scRNA-seq data to convert the base GRN into context-dependent GRN configurations for each defined cell cluster. Removal of inactive connections refines the base GRN structure, selecting for the active edges which represent regulatory connections associated with a specific cell type or state (**Figure 1B-D**). For this process, we leverage regularized machine learning regression models (Camacho et al., 2018), primarily to select active regulatory genes and to obtain their connection strength. CellOracle builds a machine learning model that predicts target gene expression from the expression levels of the regulatory genes identified in the prior base GRN refinement step. After fitting models to sample data, CellOracle extracts gene-gene connection information by analyzing model variables. With these values, CellOracle prunes insignificant or weak connections, resulting in a cell-type/state-specific GRN configuration.

CellOracle utilizes either Bayesian Ridge or Bagging Ridge models (Marbach et al., 2012; Tipping, 2001), depending on the context. These simple linear regularized machine learning models offer several advantages: (i) Regularization not only helps distinguish true factors from random, non-necessary variables but also reduces overfitting, even with a small number of observations. (ii) A linear model supports the fitting of results in an interpretable manner. Although linear models generally do not perform well with mixed complex data representing multiple regulatory states, CellOracle overcomes this limitation by inferring GRN configurations from homogenous cell types and states, based on transcriptional similarity. (iii) Rather than returning a single coefficient value for fitting results, a Bayesian/Bagging Ridge model provides a distribution of coefficient values (**Figure 1D**). This information reveals the confidence of gene-gene connections, enabling the pruning of weak or insignificant connections within the network. (iv) The linear model enables the simulation of perturbations, explained below. Together, these features position CellOracle as a unique tool to facilitate GRN inference by leveraging multiple single-cell genomic technologies (scRNA-seq and scATAC-seq), machine learning tools, and prior biological knowledge of TF-DNA interactions.

### CellOracle *in silico* simulation of transcription factor perturbations

A second key element of CellOracle is the provision of mechanistic insight into cell fate decision making by simulating the effects of TF perturbations based on the inferred GRN configurations (**Figure 1E**). In essence, CellOracle predicts how cell identity/state shifts upon a change in expression levels of a specific regulatory gene, potentially revealing how a TF determines cell identity. The simulation uses a GRN configuration to extrapolate/interpolate gene expression values, bypassing the requirement for experimental perturbation or training data.

CellOracle *in silico* perturbation consists of three steps: 1) The first step involves signal propagation simulation using a cell/state-specific GRN configuration; since CellOracle uses a linear machine learning model for GRN inference, this enables the simulation of target gene expression changes in response to changes in regulatory gene expression, regardless of other unknown factors or random errors (**Figure 1E**). By repeating this calculation n times, CellOracle simulates the n^th^ indirect effect, enabling the estimation of broad downstream effects resulting from the perturbation of a single TF. As a result, this calculates the global transcriptional effect of the perturbation; hence, we can estimate the direction of the cell identity/state transition. 2) The simulated values are compared with the gene expression of local neighbors to estimate cell identity/state transition probability. 3) CellOracle then creates a transition trajectory graph based on these transition probabilities, projecting the predicted identity of these cells upon perturbation of a candidate TF. These calculations are analogous to RNA velocity analysis, which uses RNA splicing information to infer cell state transitions (La Manno et al., 2018). Here, we adapt this method to enable visualization of CellOracle simulation, generating the results with GRN signal propagation, rather than transcript splicing information. The overall CellOracle pipeline is shown in **Figure 1F**.

### Validation of CellOracle GRN inference with ChIP-seq data

A core design element of CellOracle is the identification of context-dependent connections between a TF and its targets. Such connections are established predominantly through ‘active’ promoters/enhancers. To validate CellOracle in this context, we leveraged publicly available liver ChIP-seq datasets (**Table S1**) for histone modifications (H3K4me3 marking active promoters and H3K27ac marking active enhancers) to identify active regulatory elements. We compared inferred scores of the promoter/enhancer regions (**Figure 2A**, left) to the corresponding ChIP-seq datasets (**Figure 2A**, right). Here, we use the base promoter/enhancer information generated from the mouse scATAC-seq atlas (Cusanovich et al., 2018) for CellOracle GRN inference. Because intergenic connections are dependent on cell type and context, we matched scRNA-seq and ChIP-seq datasets across the same tissues, to enable comparisons between CellOracle and ChIP-seq. We observe significantly higher CellOracle scores for H3K4me3- and H3K27ac-positive peaks, relative to negative peaks (*P* < 0.001). To quantitatively evaluate the performance of CellOracle inference, we calculated the receiver operating characteristics (ROC) for each of these results. The area under the curve (AUC) scores in the ROC analysis were consistently high (ranging from 0.8 to 0.87; **Figure 2B, C**). We also observe a similar enrichment of CellOracle scores across active promoters and enhancers in embryonic stem cells (ESCs) (**Figure S1A-B**). Together, this epigenetic mark-based validation indicates that CellOracle accurately distinguishes the state of regulatory elements through the machine learning fitting process.

**Figure 2.**
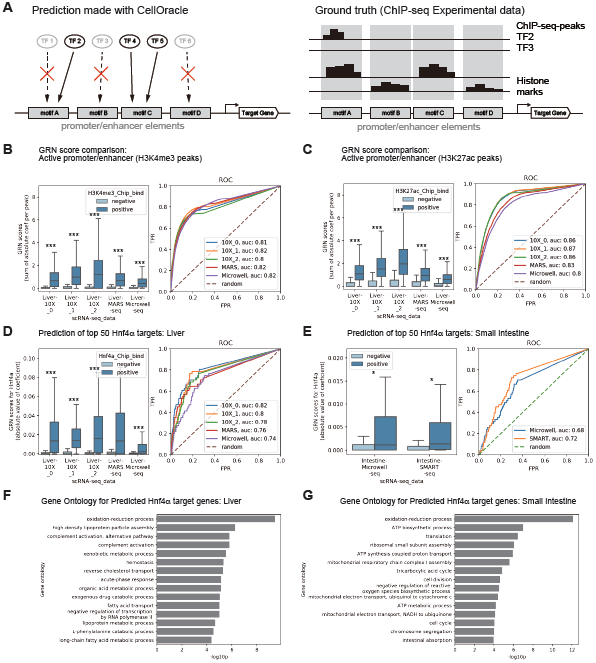
Validation of CellOracle. **(A)** CellOracle’s GRN configuration inference results were validated using existing ChIP-seq data as TF-target gene connection and active regulatory region ground truth; in this analysis, we compared inferred scores of the promoter/enhancer regions (left) to the corresponding ChIP-seq datasets (right). In addition, for a given TF and tissue, CellOracle TF-target gene scores (left) were compared to ChIP-seq peaks of the same DNA sequence (right). The existing scRNA-seq, from several platforms (10x Chromium, MARS-seq, Microwell-seq), and ChIP-seq datasets used in this analysis are detailed in **Table S1. (B-C)** Comparison of inferred promoter/enhancer element activity between positive and negative ChIP-seq peaks for active histone marks, H3K4me3 **(B)**, and H3K27ac **(C)**. The y-axis of the box plot is the inferred GRN score for each DNA peak. The score is defined as the sum of the absolute value of the GRN connection at the peak. (Box plots, dark blue: positive peaks; light blue: negative peaks; Outliers, determined by the interquartile outlier rule, are not shown; *** = *P* < 0.001). The score is defined as the sum of the absolute value of the GRN connection at the peak. ROC curves show predictive scores when we predict H3K4me3, and H3K27ac peaks based on the GRN score. **(D)** Comparison between inferred Hnf4α target genes and Hnf4α ChIP-seq experimental data for the liver and **(E)** small intestine. We selected the top 50 Hnf4α target genes, and all other genes on the basis of their promoter Hnf4α ChIP-seq peak scores. Using several existing liver and intestine scRNA-seq, CellOracle analysis was used to infer the strength Hnf4α-gene connections for these two groups (Box plots, dark blue: Hnf4α targets; light blue: non-targets; Outliers, determined by the interquartile outlier rule, are not shown; * = *P* < 0.05, *** = *P* < 0.001). The ROC curves show predictive scores when we predict the top 50 Hnf4α target genes based on the strength of the CellOracle TF-target gene connectivity scores. TPR: True Positive Rate; FPR: False Positive Rate. **(F)** Gene ontology analysis for the predicted Hnf4α target genes in the liver and **(G)** the small intestine.

Next, to validate CellOracle GRN inference, we used multiple publicly available ChIP-seq datasets (**Table S1**) to serve as ground truth physical TF-DNA interactions. To test the efficacy of CellOracle to infer cell-type/state-specific TF-gene connections, we focused on Hnf4α; a well-characterized TF redeployed in different developmental and physiological contexts (Garrison et al., 2006; Li et al., 2000; Parviz et al., 2003). Using several existing liver ChIP-seq datasets (**Table S1**), we identified the top 50 Hnf4α target genes, based on the rank of MACS2 binding scores. **Figure 2D** shows that these genes receive significantly higher Hnf4α target scores, based on CellOracle GRN inference, relative to all other genes (*P* < 0.001). These results are consistent using data collected across several different scRNA-seq platforms. We calculated the ROC curve for each of these results. Although the mean of the scores varied depending on the data and scRNA-seq method (**Figure 2D**, left panel, x-axis), the area under the curve (AUC) scores in the ROC analysis were consistently high (ranging from 0.7 to 0.8; **Figure 2D**, right panel), demonstrating the robustness of CellOracle’s GRN inference. Additionally, we benchmarked CellOracle against a current GRN inference method, GENIE3 (Huynh-Thu et al., 2010), that has recently been demonstrated to outperform other approaches (Pratapa et al., 2020). **Figure S1C-E** shows that CellOracle outperforms GENIE3 for inference of Hnf4α target genes from liver scRNA-seq data. We note here that use of a specific liver scATAC-dataset further enhances the performance of CellOracle. Finally, comparisons of Oct4/Pou5f1, Sox2, Klf4, Myc, and Nanog binding (from ChIP-seq) and inference (from CellOracle) from ESC datasets provide further validation of our approach (**Figure S1F**).

We repeated this analysis using small intestine ChIP-seq datasets, further validating CellOracle Hnf4α target gene inference (**Figure 2E**). To evaluate the specificity of CellOracle inference in the liver and intestine, we used gene ontology (GO) analysis to assess the tissue-specific functional characteristics of inferred target genes. GO terms for inferred Hnf4α target genes in the liver included well-characterized hepatic functions such as redox processes, high-density lipoprotein particle assembly, xenobiotic metabolic processes, and fatty acid transport (**Figure 2F**). In contrast, GO analysis of inferred Hnf4α targets in the small intestine includes terms related to intestinal function and the high turnover of this tissue (**Figure 2G**). Together, these results demonstrate the efficacy of CellOracle to infer cell-type/state-specific GRN configurations with a high level of sensitivity and specificity.

### Application of CellOracle to infer GRN configurations in hematopoiesis

To further assess the performance and utility of CellOracle, we applied it to a well-characterized, gold-standard model of cell differentiation: hematopoiesis (Orkin and Zon, 2008). We aimed to reproduce the reported activity of TFs regulating mouse hematopoiesis by applying CellOracle to a scRNA-seq atlas of myeloid progenitor populations (Paul et al., 2015). Dimensionality reduction using a force-directed embedding algorithm, and Louvain clustering of this 2,730 cell dataset reproduces the expected hierarchical, continuous, and branching differentiation trajectory (**Figure 3A**). Cell clusters were manually annotated, based on marker gene expression, and according to the original annotation from Paul et al., 2015. This clustering revealed the expected myeloid subpopulations, corresponding to progenitors differentiating toward erythrocytes, megakaryocytes, dendritic cells, monocytes, neutrophils, basophils, and eosinophils (**Figure 3B-D**). To analyze the network dynamics across this differentiation process, we inferred GRN configurations for each of the 24 myeloid clusters identified.

**Figure 3.**
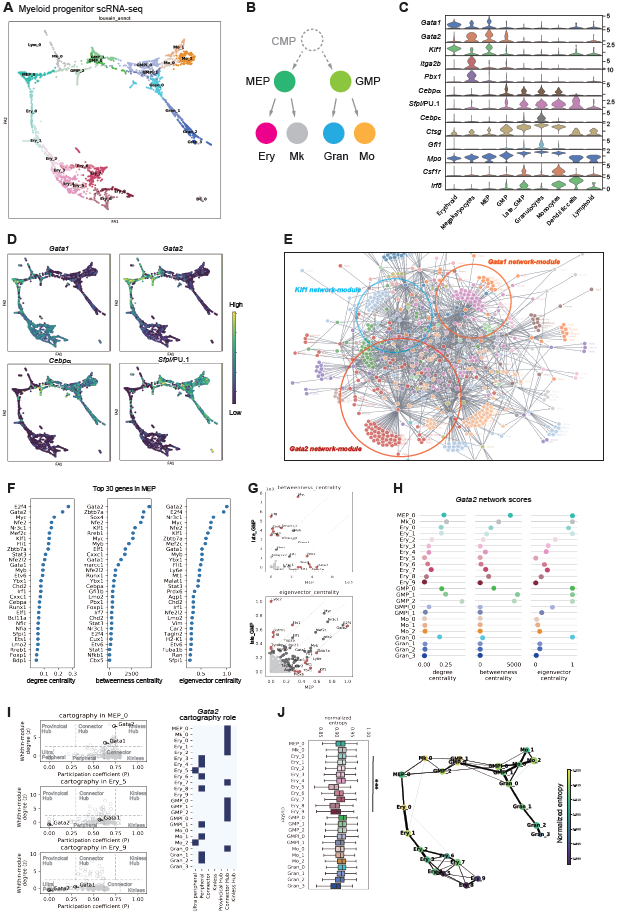
Application of CellOracle to assess GRN dynamics of hematopoiesis. **(A)** Partition-based graph abstraction (PAGA) (Wolf et al., 2019) analysis of 2,730 myeloid progenitor cells from (Paul et al., 2015). This analysis reveals 24 major clusters and reproduces the known differentiation according to a hierarchical, continuous, and branching trajectory, which is summarized as the differentiation model in **(B)** where a ‘hypothetical’ common myeloid progenitor (CMP) is shown with a dashed line. **(C)** Violin plots of marker gene expression defining each cell type in **(A). (D)** Projection of normalized and log-transformed gene expression onto the force-directed graph in **(A). (E)** Network graph of the GRN configuration for MEP cluster 0. The network edges were filtered based on p-value and strength (see methods). The graph shows the top 1,500 edges. Genes are colored according to network module membership. **(F)** Network score visualization of the MEP_0 GRN configuration centrality scores. An MEP_0 cluster-specific GRN configuration was constructed and measures of network centrality were calculated after quality checking and filtering (see methods). The top 30-scoring genes in degree centrality, betweenness centrality, and eigenvector centrality are shown here. **(G)** Scatter plots visualizing network centrality relationships between two distinct GRN configurations: late_GMP and MEP. **(H)** Analysis of *Gata2* network score dynamics across all clusters. **(I)** Gene cartography analysis. Left panels: Gene cartography scatter plots, highlighting the connectivity of *Gata2* and *Gata1* across selected clusters representing erythropoiesis. Right panel: a summary of *Gata2* gene cartography analysis, revealing its central role as a connector hub in MEPs. **(J)** Left panel: Box plot showing the distribution of network entropy scores for each cluster. *** = *P* < 0.001, Wilcoxon Test. Right panel: Projection of network entropy and degree centrality scores (median values) onto the PAGA graph.

CellOracle output initially yielded a complex network of inferred TF-to-gene connections for each cluster. To assess the properties of these networks, we first examined network degree distribution, a fundamental measure of network structure. This analysis revealed that CellOracle-inferred GRN configurations resemble a scale-free network, whose degree distribution follows a power law (**Figure S2A**). This configuration is characteristic of biological networks, defined by the presence of hub nodes serving as bridges between small degree nodes, supporting a robust network structure, in contrast to random networks (Klein et al., 2012). Many hub nodes are visible within CellOracle-inferred GRN configurations; for example, for the configuration inferred for the megakaryocyte erythrocyte progenitor (MEP) cluster, network modules and hubs can be distinguished (**Figure 3E**). The genes corresponding to these hubs, *Gata2, Gata1*, and *Klf1*, are TFs central to progenitor cell maintenance and erythrocyte differentiation (Fujiwara et al., 1996; Miller and Bieker, 1993; Tsai et al., 1994).

Visual interrogation of network modules and hubs does not represent a practical or systematic strategy by which to identify core regulators of a biological process. Therefore, to interrogate the role of individual TFs within the network, we leveraged principles of graph theory to identify critical nodes, defined by different measures of centrality. For example, degree centrality is the most straightforward measure, reporting how many edges are connected to a node (in this case, how many genes a given TF connects to) (Klein et al., 2012). Here, degree centrality scoring in the MEP_0 cluster GRN configuration successfully recognized key TFs associated with erythrocyte differentiation; *Gata1, Gata2*, and *Klf1*, as above, in addition to *Nfe2* (Andrews et al., 1993) and *Ztbt7a* (Norton et al., 2017) (**Figure 3F; Figure S2B**). However, degree centrality can often highlight TFs controlling relatively broad cellular functions, such as cell cycle and survival. Thus, we also surveyed alternate measures of network connectivity, focusing on betweenness centrality and eigenvector centrality; genes with high betweenness are essential for the transfer of information within a network, whereas genes with high eigenvector centrality scores have the most substantial influence in terms of their connections to other well-connected genes (see methods).

We compared betweenness centrality and eigenvector centrality scores across specific myeloid cluster GRN configurations, enabling the resolution of identity-specific network structures from ubiquitous network structures. For example, within the late granulocyte-monocyte progenitor (GMP) and MEP GRN configurations, *Myc*, and *Myb*, representing TFs broadly involved in the regulation of hematopoiesis (Delgado and León, 2010; Wang et al., 2018), score highly across centrality measures (**Figure 3G**). Conversely, TFs such as *Irf8, Gata2, Gata1, and Klf1*, receive variable scores between GRN configurations. Indeed, these are known regulators of hematopoietic cell fate decisions; *Gata2, Gata1*, and *Klf1* promote cell differentiation down the erythroid lineage (Fujiwara et al., 1996; Miller and Bieker, 1993; Tsai et al., 1994). Whereas previous reports suggest that *Irf8* regulates myeloid lineage commitment to monocyte and dendritic cell (DC) lineages (Becker et al., 2012); the late GMP cluster under study here associates with a DC fate (Paul et al., 2015). CellOracle successfully identified these TFs central to hematopoiesis, distinguishing those factors controlling cell identity from factors regulating ubiquitous cell processes.

### Mapping network reconfiguration during hematopoiesis

Next, we analyzed how network connectivity changes during cell differentiation. We focused specifically on Gata2, a central regulator of progenitor maintenance and erythroid differentiation. *Gata2* is predominantly expressed in hematopoietic stem and progenitor cells and is an essential factor in the maintenance of these compartments (Tsai and Orkin, 1997; Tsai et al., 1994). In the initial phase of erythroid differentiation, GATA2 induces *Gata1* expression, followed by ‘GATA factor switching’; expressed GATA1 displaces GATA2 at the *Gata2* gene to suppress its expression, driving erythropoiesis (Grass et al., 2003; Moriguchi and Yamamoto, 2014). Indeed, in the MEP cluster, *Gata1* and *Gata2* are co-expressed (**Figure 3C, D**), placing this cluster closer to the start of erythroid differentiation. In this early stage of differentiation, *Gata2* exhibits relatively high scores, across all measures of centrality, with these scores decreasing along the erythroid differentiation trajectory (**Figure 3H**). This transition is accompanied by a loss of *Gata2* expression, whereas *Gata1* expression is maintained, in line with GATA switching (**Figure 3C, D**). *Gata2* is also highly connected in GMPs, in line with previous reports of GMP-specific defects in *Gata2+/−* mice (Rodrigues et al., 2008). In contrast to the erythroid lineage, we observe a drastic decrease in *Gata2* network scores along the GMP differentiation trajectory (**Figure 3H**).

To further interpret *Gata2* connectivity, we performed network cartography analysis. This method uses the topology of the network to classify genes into several groups based on intra- and inter-module connections of the GRN configuration, revealing putative roles for candidate TFs (**Figure 3I**; **Figure S3**). Using this approach, *Gata2* was classified as a connector hub in the MEP cluster (**Figure 3I**), suggesting that *Gata2* employs multiple regulatory mechanisms by linking genes both within and between modules, acting as an important connector. In contrast to this progenitor cluster, *Gata2* cartography analysis in the differentiated erythroid cell clusters shows that *Gata2* loses regulatory connections as erythroid differentiation progresses (**Figure 3I**; **Figure S3**). Together, these results agree with the known role of *Gata2* as a master regulator in hematopoiesis, demonstrating the sensitivity of CellOracle to characterize GRN reconfiguration during cell differentiation.

Finally, we analyzed network entropy scores to gain insight into the global features of GRN dynamics. Network entropy is affected by the activation of specific signaling pathways; thus, the entropy score negatively correlates with differentiation state (Banerji et al., 2013; Teschendorff and Enver, 2017). **Figure 3J** shows network entropy distribution, calculated from the distribution of network edge strength. As expected, the network entropy score significantly decreases as cells differentiate (Ery_0 vs. Ery_9, *P* < 0.001, Wilcoxon Test), implying that the inferred GRN configurations reflect regulatory pathway transitions in response to differentiation.

### Simulating the effects of TF knockout on myeloid cell identity

CellOracle GRN inference and network analysis successfully identified central TF regulators of hematopoiesis. To demonstrate how the role of these TFs can be further investigated and prioritized for additional analysis using CellOracle, we performed *in silico* perturbation for *Gata1* and *Sfpi1* (encoding PU.1). These two factors mutually antagonize each other in the erythroid vs. myeloid fate decision to generate MEPs and GMPs, respectively (Rekhtman et al., 1999; Zhang et al., 1999). PU.1 expression is high in GMPs, directing commitment to the neutrophil and monocyte lineages (Back et al., 2005; Nutt et al., 2005). Conversely, Gata1 promotes erythroid differentiation (Fujiwara et al., 1996). Using CellOracle-based simulation of *Gata1* and *Sfpi1/*PU.1 perturbation, we aimed to recapitulate the lineage switch between MEP and GMP identities.

Using the 24 hematopoietic GRN configurations inferred by CellOracle, we simulated *Gata1* knockout signal propagation, enabling the future gene expression and hence the direction of cell identity transitions to be predicted, at single-cell resolution (**Figure 4A**). This simulation predicts a clear shift of MEP cell identity toward a GMP signature, as would be expected following *Gata1* knockout (**Figure 4B**). To predict changes in cell-type composition, in **Figure 4C**, the transition probability of an individual cell (red diamond), after *Gata1* knockout, is projected onto the plot. A Markov chain simulation is performed for this individual cell to simulate a change in identity/state. We then simulated cell transitions for all cells in the population, predicting an increase in cell density toward the GMP cluster and early erythroid branch upon *Gata1* knockout (**Figure 4D**). This increase in erythroid precursors is in agreement with the observed arrest of gene-disrupted GATA1 null cells at the proerythroblast stage *in vitro* (Fujiwara et al., 1996). In addition, breaking down the direction of these predicted cell transitions by cluster reveals a shift in cell identity from granulocytes to late GMPs (**Figure 4E**). In contrast to these results, simulation of *Gata1* overexpression predicts a transition toward MEP identities, in addition to granulocytes (**Figure S4A-E**). Finally, our simulation of *Sfpi1/*PU.1 knockout predicts the opposite effects, compared to the *Gata1* knockout simulation results, as expected for these mutually antagonizing factors (**Figure 4F-J**).

**Figure 4.**
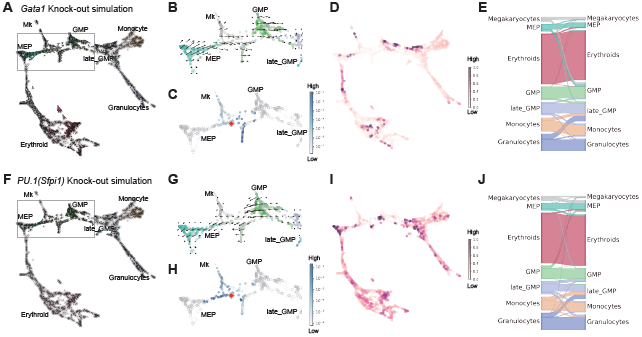
CellOracle simulation of *Gata1* and *Sfpi1*/PU.1 knockout during hematopoiesis. CellOracle prediction of cell state transitions in *Gata1* knockout **(A-E)** and *Sfpi1*/PU.1 knockout simulations **(F-J). (A-B/F-G)** Projection of cell state transition vectors for each cell on the force-directed graph. **(B/G)** Magnified area of the graph surrounded by the light-gray rectangle in **(A)**, with projection of the local average of the transition vector calculated per grid point. **(C/H)** Magnified area of the graph in **(A/F)**, with projection of cell transition probability defined by the similarity of gene expression between simulated cells and other pre-existing cells. This plot shows the transition probability score for one specific cell (representing a mid-point between MEP and GMP lineages), highlighted by a red diamond. After perturbation (simulated loss of *Gata1, or Sfpi1*/PU.1 expression), this cell is predicted to adopt a similar gene expression profile to the cells colored in blue, denoting an increasing transition probability. **(D/I)** A Markov simulation (number of simulation steps = 50) was performed with the transition probability for all cells in the population, resulting in a cell density estimation following gene knockout. The magenta color denotes a higher cell density, and white denotes a lower cell density. **(E/J)** A Sankey diagram showing cell transitions between different cell types in the Markov simulation.

To further validate CellOracle simulation with additional TFs, we performed *in silico* knockouts of *Cebpα* and *Cebpε.* Experimental knockout previously demonstrated that *Cebpα* is necessary for initial GMP differentiation, where its loss leads to a drastic decrease in differentiated myeloid cells, accompanied by an increase in erythroid progenitor cell numbers (Paul et al., 2015). CellOracle perturbation simulation reproduces these phenotypes (**Figure S4F-O**), where we predict a clear transition from the GMP to MEP lineage upon simulated *Cebpα* knockout (**Figure S4F-J**). In the context of *Cebpε*, its experimental knockout blocks granulocyte differentiation, leading to an increase of GMPs (Paul et al., 2015). Again, CellOracle’s *Cebpε* knockout simulation recapitulates these experimental findings: cell differentiation arrests during granulocyte differentiation, as expected, resulting in an increase in cell density within the late GMP cluster (**Figure S4K-O**).

Together, the results of our CellOracle simulation of *Gata1, Sfpi1/*PU.1, *Cebpα* and *Cebpε* perturbations successfully recapitulate previous experimental perturbations of these factors. These simulations demonstrate the capacity of CellOracle to elucidate complex biological mechanisms, such as the molecular switches of lineage determination. In addition to these major lineage determinants, CellOracle analysis also enabled the intuitive interpretation of additional cell identity transitions predicted via *in silico* perturbation. For example, simulation of *Gata1* knockout cell transitions reveals a shift from granulocyte to late GMP cell identity, implying a positive role for Gata1 in granulocyte differentiation. Our *Gata1* overexpression simulation supports this conclusion. Supporting our observations here, in addition to the primary role of Gata1 to promote MEP lineage differentiation, it was reported that Gata1 is necessary for the terminal maturation of granulocytes, basophils specifically (Nei et al., 2013). Here, CellOracle successfully identified the multiple roles of Gata1 in hematopoiesis; Gata1 is central to both erythropoiesis, and granulocyte terminal differentiation. These results demonstrate that CellOracle simulation can reproduce the diverse, context-specific roles of TFs.

### Mapping network configuration changes during cell fate reprogramming

We have so far demonstrated the efficacy of CellOracle to recapitulate known biology, successfully identifying TFs central to cell differentiation and correctly predicting the outcome of their perturbation on cell identity. Next, we applied CellOracle to cells undergoing reprogramming, aiming to reveal new mechanistic insights into the TF-mediated conversion of cell identity. Here, we focus on direct lineage reprogramming, which intends to convert one mature cell type directly into another, bypassing pluripotent or progenitor states (Cohen and Melton, 2011; Vierbuchen and Wernig, 2011). However, such reprogramming strategies are typically inefficient and fail to fully recapitulate target cell identity (Morris and Daley, 2013). Thus, reprogramming represents an ideal application for CellOracle to identify factors that can enhance the efficiency and fidelity of cell fate conversion.

We have previously investigated the direct lineage reprogramming of mouse embryonic fibroblasts (MEFs) to induced endoderm progenitors (iEPs), generated via the forced expression of two TFs: Hnf4α and Foxa1 (**Figure 5A;** (Biddy et al., 2018; Morris et al., 2014)). The generation of iEPs represents a prototypical lineage reprogramming protocol, which, like most conversion strategies, is inefficient and lacks fidelity. The resulting cells, initially described as hepatocyte-like cells, can functionally engraft the liver (Sekiya and Suzuki, 2011). However, we previously demonstrated that these cells also harbor intestinal identity and can functionally engraft the colon in a mouse model of acute colitis, prompting their re-designation as iEPs (Guo et al., 2019; Morris et al., 2014). Building on these studies, our recent single-cell lineage tracing of this protocol revealed two distinct trajectories arising during MEF to iEP conversion: one to a successfully reprogrammed state, and one to a dead-end state, where cells fail to fully convert to iEPs (Biddy et al., 2018). Although we had identified factors to improve the efficiency of reprogramming, mechanisms of cell fate conversion from the viewpoint of GRN dynamics remain unknown. Here, we deploy CellOracle to interrogate GRN reconfiguration during MEF to iEP conversion, leveraging this knowledge to improve reprogramming efficiency.

**Figure 5.**
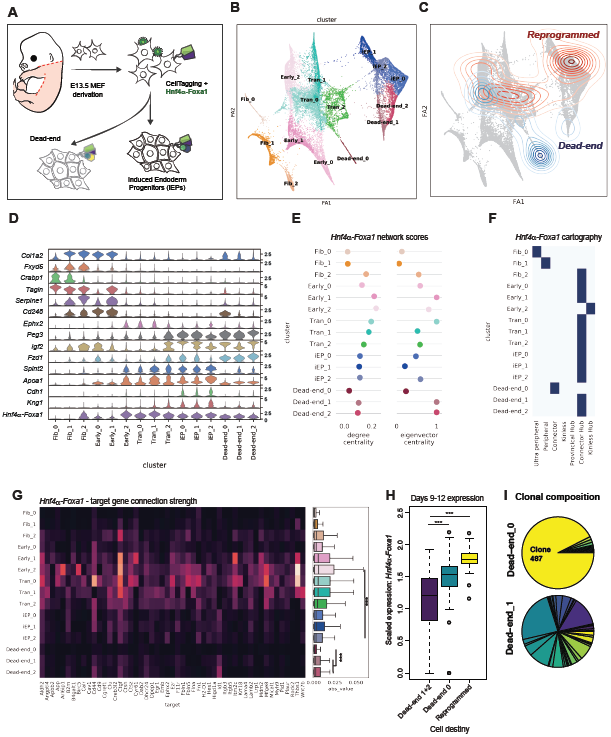
Application of CellOracle to assess GRN dynamics direct lineage reprogramming. **(A)** Schematic of fibroblast to iEP reprogramming, driven by overexpression of Hnf4α and Foxa1. Our previous CellTag lineage tracing revealed two conversion trajectories; one yielding successfully converted iEPs, and a path leading to a ‘dead-end’ state (Biddy et al., 2018). scRNA-seq data of this 28-day reprogramming timecourse was analyzed with CellOracle to investigate GRN dynamics over the cell reprogramming process. **(B)** Force-directed graph of fibroblast to iEP reprogramming: from Louvain clustering, 15 groups of cells were distinguished. Cell clusters were annotated manually, using marker gene expression, and grouped into five cell types; Fibroblasts, Early_Transition, Transition, Dead-end, and Reprogrammed iEPs. **(C)** Projection of cell lineage information, collected using the CellTag lineage tracing method (Biddy et al., 2018), onto the force-directed graph. Kernel density estimation of CellTag lineage information: cells belonging to the dead-end trajectory are visualized in blue, while cells on the successfully reprogrammed trajectory are shown in red. **(D)** Violin plots of select marker gene expression for each cluster shown in **(B). (E-G)** Network analysis of the reprogramming transgene, *Hnf4α-Foxa1*. **(E)** Network score dynamics inferred by CellOracle: CellOracle was applied to iEP reprogramming scRNA-seq data to reconstruct cluster-specific GRN configurations. Degree centrality, betweenness centrality, eigenvector centrality of *Hnf4α-Foxa1* for each cluster are shown here. **(F)** Network cartography terms of *Hnf4α-Foxa1* for each cluster. **(G)** The strength of network edges between *Hnf4α-Foxa1* and its target genes, visualized as a heatmap (left panel), and plotted as a boxplot (right panel). **(H)** Transgene expression levels (at Days 9-12) in cells destined to Dead-end clusters 1+2, Dead-end cluster 0 and reprogrammed iEP clusters (*P* < 0.01, *t*-test, one-sided). **(I)** Pie charts depicting the clonal composition of Dead-end cluster 0 and Dead-end cluster 1. Clone and trajectory information in **(H, I)** is derived from our previous CellTagging study (Biddy et al., 2018).

Our previously published MEF to iEP reprogramming scRNA-seq dataset consists of eight time points, collected over 28 days (*n* = 27,663 cells) (Biddy et al., 2018). Following preprocessing, dimensionality reduction, and clustering, we manually annotated 15 clusters using partition-based graph abstraction (PAGA; (Wolf et al., 2019)), and marker gene expression (**Figure 5B; Figure S5**). In agreement with our previous analyses, fibroblasts gradually convert to iEPs, progressing through the ‘early-transition’ and ‘transition’ phases. After successfully initiating conversion, these cells then diverge down one of two trajectories: one leading to a successfully reprogrammed state (**Figure 5C**: red cells), and one to a dead-end state (**Figure 5C**, blue cells). Cells along both paths proliferate extensively, whereas unconverted fibroblasts senesce. From our previous CellTagging-based (Kong et al., 2020a) lineage analysis, these trajectories can be tracked to earlier transition stages, revealing early differences between cells on these paths that determine reprogramming outcome (**Figure 5C**; (Biddy et al., 2018)). Cells within the dead-end clusters exhibit distinct gene expression patterns, relative to fully reprogrammed iEP clusters. Notably, dead-end cells only weakly express iEP markers, *Cdh1*, and *Apoa1*, accompanied by higher expression levels of fibroblast marker genes, such as *Col1a2* (**Figure 5D; Figure S5C**). Using CellOracle, we inferred GRN configurations for each cluster, calculating network connectivity scores to analyze GRN dynamics during lineage reprogramming.

### *Hnf4α-Foxa1* transgene network configurations reveal different modes of reprogramming failure

Reprogramming to iEPs is driven by Hnf4α and Foxa1 TFs, delivered via retrovirus and expressed as a bicistronic transcript, *Hnf4α-t2a-Foxa1*, for consistent reprogramming factor stoichiometry. We initially focused on the network configuration associated with *Hnf4α-Foxa1*, where this transgene receives high degree centrality and eigenvector centrality scores in the early phases of lineage conversion, gradually decreasing as reprogramming progresses (**Figure 5E**). Hnf4α and Foxa1 receive a combined score in these analyses since they express as a single transcript that produces two independent factors via 2A-peptide-mediated cleavage (Liu et al., 2017). In agreement with a central role for these transgenes early in reprogramming, network cartography analysis classified *Hnf4α-Foxa1* as a prominent “connector hub” in the early_2 cluster network configuration (**Figure 5F**). Indeed, network strength scores show significantly stronger connections of *Hnf4α-Foxa1* to its inferred target genes in the early stages of reprogramming, followed by decreasing connection strength in later conversion stages (Early_2 vs. iEP_2: *P* < 0.001, Wilcoxon Test; **Figure 5G**). Together, these analyses reveal that both *Hnf4α-Foxa1* network configuration connectivity and strength peak in early reprogramming phases. Our previous observations that reprogramming outcome is determined shortly after initiation of lineage conversion support a crucial role for these transgenes early in conversion (Biddy et al., 2018), and is in line with the role of Foxa1 as a pioneer factor that engages with silent, unmarked chromatin to initiate transcriptional changes resulting in the conversion of cell identity (Iwafuchi-Doi and Zaret, 2016).

Next, we analyzed the *Hnf4α-Foxa1* network configuration in later stages of conversion, following bifurcation into reprogrammed and dead-end trajectories at the 21-day time point (**Figure 5B, C, Figure S5B**). In the reprogrammed clusters (iEP_0, iEP_1, iEP_2) and dead-end clusters (Dead-end_1, Dead-end_2), *Hnf4α-Foxa1* retains its classification as connector hub, although network strength scores are significantly weaker, for both conversion outcomes (Early reprogramming vs. late reprogramming; *P* < 0.001, Wilcoxon Test). The reprogrammed clusters exhibit stronger network connectivity scores, relative to the dead-end clusters (Dead-end vs. iEP; *P* < 0.001, Wilcoxon Test). In addition, the levels of *Hnf4α-Foxa1* expression differ between outcomes, with significantly higher transgene expression observed in reprogrammed clusters (*P* < 0.001, *t-*test, one-sided; **Figure 5D**). To investigate this further, we leveraged our existing CellTag lineage tracing data to probe *Hnf4α-Foxa1* expression levels in cell ancestors at earlier stages of reprogramming, before trajectory bifurcation. Cells destined to the dead-end outcome express significantly lower levels of *Hnf4α-Foxa1* (*P* < 0.001, *t-*test, one-sided; **Figure 5H**). Furthermore, CellOracle GRN inference reveals that the networks of reprogrammed vs. dead-end cells are already configured differently at early stages (**Figure S6A-C;** Days 6-15, n = 51 dead-end cells, 42 reprogrammed cells). Notably, degree centrality and eigenvector centrality scores for *Zeb1* in the early dead-end trajectory are higher, relative to cells destined to reprogram successfully (**Figure S6C**). We also observe increased *Zeb1* expression in dead-end clusters at day 28 (**Figure S6D;** *P* < 0.001, permutation test, one-sided). Zeb1 is a TF associated with the promotion of epithelial to mesenchymal transition (Liu et al., 2008); thus the induction and expression of this factor may account for the observed persistence of fibroblast marker gene expression, absence of iEP marker expression, and failure to complete a mesenchymal to epithelial transition (MET) along the dead-end (**Figure 5D**). Indeed, *Zeb1* knockout simulation predicts that loss of this TF shifts cells from the dead-end, into a reprogrammed state (**Figure S6E**).

In addition to the above dead-end clusters (1 and 2), our new clustering presented in this study reveals a third, distinctive dead-end trajectory (Dead-end_0). From gene expression, cells within this cluster appear to only weakly initiate reprogramming, retaining robust fibroblast gene expression signatures and expressing significantly lower levels of reprogramming initiation markers such as *Apoa1* (*P* < 0.001, permutation test; **Figure 5D**). This cluster exhibits distinctive patterns in terms of transgene network connectivity; degree centrality and eigenvector centrality scores for *Hnf4α-Foxa1* in the Dead-end_0 cluster are significantly lower, relative to the other dead-end clusters (Dead-end_1 and Dead-end_2; *P* < 0.001, Wilcoxon Test; **Figure 5E**). In support of this observation, cartography analysis defines *Hnf4α-Foxa1* in the Dead-end_0 cluster as a “Connector,” indicating that this transgene has a relatively small number of connections within the module. In contrast, the other dead-end clusters receive a relatively more connected “Connector Hub” status (**Figure 5F**). Furthermore, the connection strength of *Hnf4α-Foxa1* in the dead-end_0 cluster is significantly weaker compared to the other dead-end clusters (*P* < 0.001, Wilcoxon Test; **Figure 5G**). These results indicate that *Hnf4α-Foxa1* connectivity is significantly attenuated in the dead_end_0 network configuration.

Again, we analyzed transgene levels in ancestors before their emergence onto this distinctive dead-end path. Unexpectedly, Dead-end_0-destined cells express significantly higher levels of *Hnf4α-Foxa1*, relative to Dead-end_1- and Dead-end_2-destined cells (*P* = 0.001, *t-*test, one-sided; **Figure 5H**). Further investigation revealed that the majority of the cells (93% of tracked cells) on this unique path derive from a single clone, representing a rare reprogramming event that we have captured due to clonal expansion (**Figure 5I**). Thus, although the cell giving rise to this clone received sufficient levels of *Hnf4α-Foxa1* to drive reprogramming, CellOracle analysis suggests that the reprogramming factors were unable to engage with the target genes required to initiate and complete lineage conversion. This possibility may be explained by *Hnf4α-Foxa1* targets being ‘locked’ in heterochromatin in this particular cell giving rise to the Dead-end_0, as has previously been suggested in other reprogramming contexts (Soufi et al., 2012). Altogether, the MEF to iEP reprogramming network analysis presented here suggests that *Hnf4α-Foxa1* function peaks at the initiation of conversion. These early, critical changes in GRN configuration determine reprogramming outcome, with dysregulation or loss of this program leading to dead-ends, where cells either do not successfully initiate or complete reprogramming. This hypothesis is consistent with our previous CellTag lineage tracing, showing the establishment of reprogramming outcomes from early stages of the conversion process (Biddy et al., 2018).

### The AP-1 transcription factor subunit Fos is central to reprogramming initiation and maintenance of iEP identity

To identify TFs with pivotal roles in reprogramming initiation, we compared network connectivity scores between cluster-specific GRN configurations. First, comparing degree centrality scores between fibroblast and early reprogramming clusters reveals that all TFs receive similar scores, except for a small number of factors (**Figure 6A**). Among these factors, *Fos* and *Zfp57* receive relatively high degree and eigenvector centrality scores, along with connector hub classification in the early reprogramming clusters (**Figure 6A-D; Figure S7A, B**). Fos is a subunit of the activator protein-1 TF (AP-1), a dimeric complex primarily containing members of the Fos and Jun factor families (Eferl and Wagner, 2003). As part of the AP-1 complex, Fos plays broad roles in proliferation, differentiation, and apoptosis, both in the context of development and tumorigenesis (Eferl and Wagner, 2003; Jochum et al., 2001; Velazquez et al., 2015). In addition, AP-1 functions to establish cell-type-specific enhancers and gene expression programs (Heinz et al., 2010; Vierbuchen et al., 2017), and to reconfigure enhancers during reprogramming to pluripotency (Knaupp et al., 2017; Madrigal and Alasoo, 2018). Zfp57 is also implicated in the control of cell identity, functioning as a transcriptional repressor (Alonso et al., 2004), maintaining repressive epigenetic modifications at imprinting control regions (Li et al., 2008; Riso et al., 2016; Shi et al., 2019). We computationally and experimentally validate a role for Zfp57 in reprogramming initiation, as shown in **Figure S7C-H**, focusing on the role of Fos in the remainder of this section.

**Figure 6.**
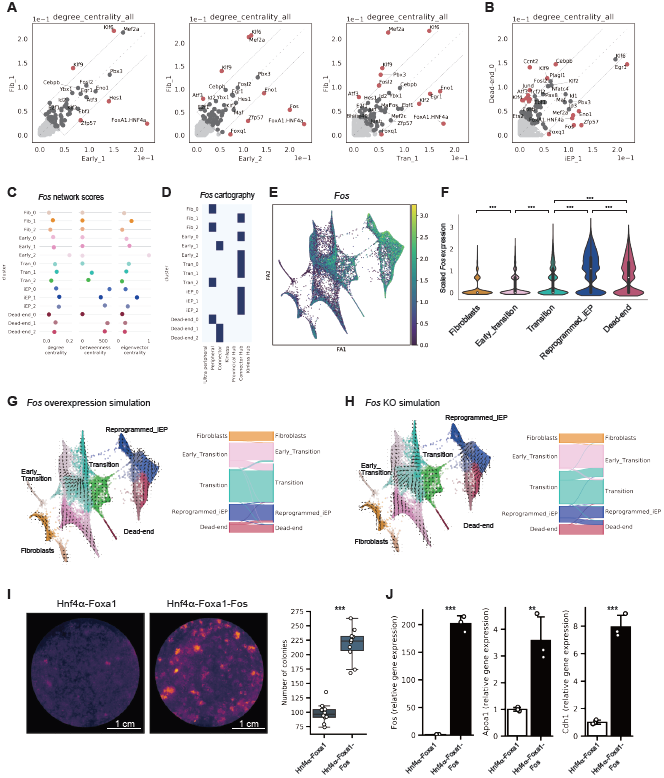
CellOracle analysis and experimental validation of Fos in iEP reprogramming initiation. **(A, B)** Scatter plots showing a comparison of degree centrality scores between specific clusters. **(A)** Comparison of degree centrality score between the Fib_1 cluster GRN configuration and the GRN configurations of other clusters in relatively early stages of reprogramming. **(B)** Comparison of degree centrality score between iEP_1 and Dead-end_0 cluster GRN configurations. **(C)** Network score dynamics inferred by CellOracle. Degree centrality, betweenness centrality, eigenvector centrality of *Fos* for each cluster are shown here. **(D)** Network cartography terms of *Fos* for each cluster. **(E)** *Fos* expression projected onto the force-directed graph of fibroblast to iEP reprogramming. **(F)** Violin plot of *Fos* expression across reprogramming stages. **(G)** *Fos* gene overexpression simulation with reprogramming GRN configurations. The left panel is the projection of simulated cell transitions onto the force-directed graph. The Sankey diagram summarizes the simulation of cell transitions between cell clusters. For the simulation of *Fos* overexpression, we set the *Fos* expression value at 1.476, which is the maximum value of *Fos* expression in the imputed gene expression matrix **(H)** *Fos* gene knockout simulation. **(I)** Colony formation assay with addition of Fos to the Hnf4α-Foxa1 reprogramming cocktail. Left panel: E-cadherin immunohistochemistry. Right panel: box plot of colony numbers (n = 6 technical replicates, 2 independent biological replicates; *** = *P* < 0.001, *t-*test, one-sided). **(J)** qPCR assay for *Fos* and iEP marker expression (*Apoa1* and *Chd1*) following addition of Fos to the Hnf4α-Foxa1 reprogramming cocktail (n = 3 technical replicates, 1 biological replicate; *** = *P* < 0.001, ** = *P* < 0.01, *t-*test, one-sided).

During MEF to iEP reprogramming, *Fos* is gradually and significantly upregulated (**Figure 6E**; *P* < 0.001, permutation test, one-sided). Several Jun AP-1 subunits are also expressed in iEPs, classifying as connectors and connector hubs across various reprogramming stages (**Figure S8A-C**). *Fos* and *Jun* are among a battery of genes reported to be upregulated in a cell-subpopulation specific manner, in response to cell dissociation-induced stress, potentially leading to experimental artifacts (van den Brink et al., 2017). Taking this report into consideration, we performed qPCR for *Fos* on dissociated and undissociated cells. This orthogonal validation confirms an 8-fold upregulation (*P* <0.01, *t-*test, one-sided) of *Fos* in iEPs, relative to MEFs, revealing no significant changes in gene expression in cells that are dissociated and lysed, versus cells lysed directly on the plate (**Figure S8D**). Furthermore, analysis of unspliced and spliced *Fos* mRNA levels reveals an accumulation of spliced *Fos* transcripts in reprogrammed cells. This observation suggests that these transcripts accumulated over time, rather than by rapid induction of expression in the five-minute cell dissociation and methanol fixation in our single-cell preparation protocol (**Figure S8E**) (La Manno et al., 2018). Taken together, based on these gene expression patterns and high network connectivity in early reprogramming, we selected *Fos* as a candidate gene playing a critical role in the initiation of iEP conversion.

To investigate a potential role for *Fos* in reprogramming initiation, we simulated overexpression signal propagation, using the MEF to iEP reprogramming GRN configurations inferred by CellOracle. Overexpression simulation for Fos predicts a major cell state shift from the early transition to transition clusters, in addition to predicting shifts in identity from the dead-end to reprogrammed clusters **(Figure 6G)**. In contrast, simulation of *Fos* knockout produces the opposite results **(Figure 6H).** Next, we experimentally validated this simulation by adding Fos to the iEP reprogramming cocktail. As expected, we see a significant increase in the number of iEP colonies formed (*P* < 0.001, *t-*test, one-sided; **Figure 6I**), increasing reprogramming efficiency more than two-fold, accompanied by significant increases in iEP marker expression (*P* < 0.001, *t-*test, one-sided; **Figure 6J**).

Turning our attention to the later stages of reprogramming, *Fos* continues to receive relatively high network scores, particularly for betweenness centrality, in the iEP GRN configurations **(Figure 6C)**. *Fos* also classifies as a Connector Hub **(Figure 6D)** in iEPs, suggesting a role for Fos in the stabilization and maintenance of the reprogrammed state. To test this hypothesis, we use CellOracle to perform knockout simulations, followed by experimental knockout validation in an established iEP cell line. Here, we leverage the ability to culture iEPs, long-term, where they retain a range of phenotypes (from fibroblast-like to iEP states; **Figure S8F**) and functional engraftment potential (Guo et al., 2019; Morris et al., 2014). Simulation of *Fos* knockout using these long-term cultured iEP GRN configurations predicts the attenuation of iEP identity upon factor knockout (**Figure 7A**). To test this prediction, we used Perturb-seq, a combination of CRISPR-Cas9 based knockout with scRNA-seq (Adamson et al., 2016). Quantitative comparison of the cell proportions between control and knockout groups confirms that fully reprogrammed iEPs regress toward an intermediate state upon *Fos* knockout, confirming a role for this factor in maintaining iEP identity (**Figure 7B**).

**Figure 7.**
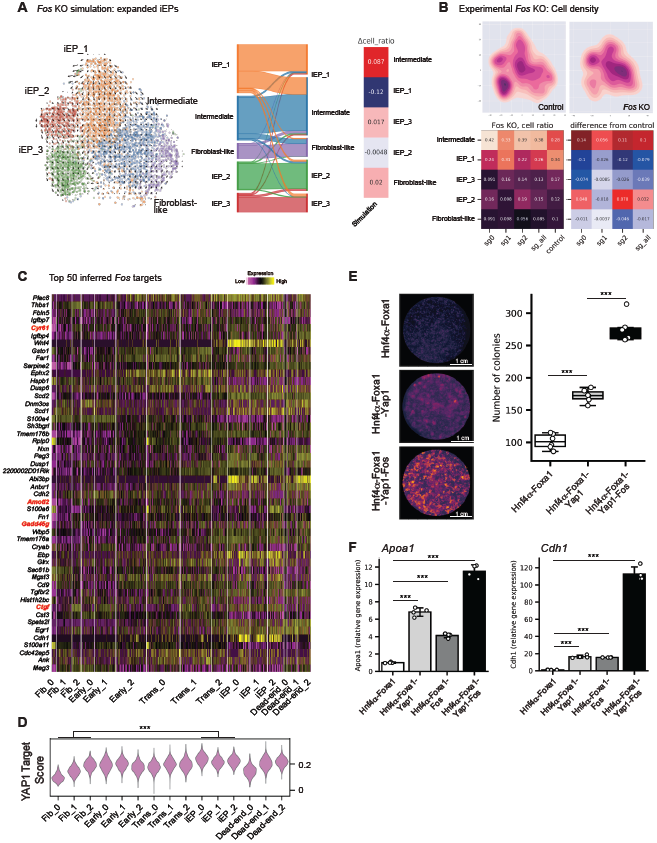
Fos is required for maintenance of cell identity; inferred Fos targets reveal a role for the Hippo signaling effector, Yap1 in reprogramming. **(A)** *Fos* gene knockout simulation in expanded, long-term cultured iEPs. **(B)** Experimental knockout of *Fos* using Perturb-seq. These experiments were performed with reprogrammed, expanded iEP cells, which express the *Cas9* gene. We designed 3 guide RNAs to target *Fos*, and transduced Cas9 iEP cells with this guide RNA lentivirus pool at low MOI (see methods). Left panels: Kernel density estimation method was applied with the t-SNE embedding to compare cell density between control guide RNAs and guide RNAs targeting *Fos*. Right panels: Quantification of changes in cell ratio following *Fos* knockout. **(C)** Heatmap of expression of the top 50 inferred *Fos* targets across all stages of reprogramming. Established targets of YAP1 are highlighted in red. **(D)** Violin plot of YAP1 target gene scores across reprogramming, which are significantly enriched as reprogramming progresses (*** = *P* < 0.001, permutation test, one-sided). **(E)** Colony formation assay with addition of Yap1 and Fos to the Hnf4α-Foxa1 reprogramming cocktail. Left panels: E-cadherin immunohistochemistry. Right panel: box plot of colony numbers (n = 6 biological replicates; *** = *P* < 0.001, *t-*test, one-sided). qPCR assay for iEP marker expression (*Apoa1* and *Chd1*) following addition of Yap1 and Fos to the Hnf4α-Foxa1 reprogramming cocktail (n = 4 biological replicates; *** = *P* < 0.001, ** = *P* < 0.01, *t-*test, one-sided).

### *Fos* target inference uncovers a role for the hippo signaling effector Yap1 in reprogramming

To gain further insight into the mechanism of how Fos regulates reprogramming, we interrogated a list of the top 50 inferred *Fos* targets across all stages of reprogramming (**Figure 7C; Table S2**). We also assembled a list of genes predicted to be downregulated following *Fos* knockout simulation for the reprogramming timecourse (**Figure S8G**). From this analysis, we noted the presence of direct targets of YAP1, a central downstream transducer of the Hippo signaling pathway (Galli et al., 2015; Ramos and Camargo, 2012; Stein et al., 2015). These targets include *Cyr61, Amotl2, Gadd45g*, and *Ctgf*. Previous links between Yap1 and Fos support these observations; for example, YAP1 is recruited to the same genomic regions as FOS, via complex formation with AP-1 (Zanconato et al., 2015). Moreover, AP-1 is required for YAP1-regulated gene expression and the liver overgrowth caused by Yap overexpression, where FOS induction contributes to the expression of YAP/TAZ downstream target genes (Koo et al., 2020).

Together, this evidence suggests that Fos may play a role in reprogramming via an AP-1-Yap1-mediated mechanism. Since Yap1 does not directly bind to DNA, we cannot deploy CellOracle here to perform network analysis or perturbation simulations. In lieu of these analyses, we turn to our rich single-cell timecourse of iEP reprogramming. Using a well-established active signature of Yap1 (Dong et al., 2007), we find significant enrichment of this signature as reprogramming progresses (**Figure 7D;** *P* < 0.001, permutation test, one-sided; **Figure S8H**). Together, these results suggest a role for the Hippo signaling component Yap1 in reprogramming, potentially effected via its interactions with Fos/AP-1. Indeed, we find that addition of Yap1 to the Hnf4a-Foxa1 significantly enhances reprogramming efficiency, where addition of both Fos and Yap1 increase colony formation by almost three-fold, accompanied by significant increases in iEP marker expression (**Figure 7E-F; Figure S8I**, *P* < 0.001, *t-*test, one-sided). Further supporting a role for Yap1 in the establishment of iEP identity are previous studies demonstrating a role for this pathway in liver regeneration (Pepe-Mooney et al., 2019; Yimlamai et al., 2014) and regeneration of the colonic epithelium (Yui et al., 2018), in line with the known potential of iEPs to functionally engraft the liver and intestine (Guo et al., 2019; Morris et al., 2014; Sekiya and Suzuki, 2011). In summary, CellOracle analysis and *in silico* prediction, in combination with experimental validation, has revealed several new factors and putative regulatory mechanisms to enhance the efficiency and fidelity of reprogramming.

## Discussion

Here, we have presented CellOracle, a machine learning-based tool to infer GRNs via the integration of different single-cell data modalities. CellOracle aims to overcome current challenges in GRN inference by using single-cell transcriptomic and chromatin accessibility profiles, integrating prior biological knowledge via regulatory sequence analysis to infer transcription factor (TF)-target gene interactions. Furthermore, we have designed CellOracle to apply inferred GRNs to the simulation of gene expression changes in response to TF perturbation. This unique feature enables inferred GRN configurations to be interrogated *in silico*, facilitating their interpretation. We benchmarked CellOracle against ground-truth TF-gene interactions and existing GRN inference tools, demonstrating its efficacy to recapitulate the diverse, context-specific roles of TFs in hematopoiesis.

We developed CellOracle to address previous limitations in GRN inference. Foremost, CellOracle uses single-cell data, enabling the deconstruction of population heterogeneity. Several platforms currently exist to reconstruct GRNs from single-cell expression data (Pratapa et al., 2020); however, the use of expression data alone can lead to poor performance (Chen and Mar, 2018) from false-positive edges forming feedforward loops (Pratapa et al., 2020). SCENIC (Single-Cell rEgulatory Network Inference and Clustering) can potentially overcome this issue by incorporating TF binding site information (Aibar et al., 2017). Here, CellOracle builds on this strategy, integrating prior knowledge on TF binding with chromatin accessibility information from scATAC-seq. The machine learning model leveraged by CellOracle integrates this multi-omic information to provide gene-gene interaction confidence scores, defining the directionality of connections and pruning weak edges to minimize false-positives. Indeed, CellOracle outperforms a currently existing best-in-class GRN inference tool (Huynh-Thu et al., 2010; Pratapa et al., 2020). Still, we acknowledge that the use of single-cell expression data may be fundamentally limited in that the reduced resolution cannot capture the gene expression variability required for reliable GRN inference. Moreover, gene expression alone may not faithfully reflect regulatory interactions. In this instance, emerging multi-omic platforms (Stuart and Satija, 2019) will prove invaluable for GRN inference, where these data modalities can be integrated into CellOracle.

A second key element of CellOracle yields mechanistic insight into cell fate decision making by using inferred GRN configurations to simulate the effects of TF perturbations. The simulation uses a GRN configuration to extrapolate/interpolate gene expression values, bypassing the requirement for experimental perturbation or training data. This functionality, together with the network connectivity scoring and cartography strategies presented here, enables intuitive interpretation of the inferred GRN configurations, allowing the prioritization of factors via *in silico* simulation, before further experimental validation. Other methods to simulate perturbations exist, but not for TFs. For example, scGen combines variational autoencoders with vector arithmetic to support ‘out-of-sample’ predictions of dose and infection response of cells (Lotfollahi et al., 2019). However this requires extensive training data for closely-related cells, contrasting to CellOracle which does not require perturbation data.

In contrast, although CellOracle-based simulation does not require TF perturbation training data to make predictions, it does require sampling of the potential cell types and states that result from a perturbation. In addition, because CellOracle uses a linear regression model, it cannot predict non-linear, combinatorial effects, limiting simulations to individual TFs. However, the simple linear regularized machine learning model deployed here does offer several advantages, reducing overfitting and delivering interpretable results. This benefit is enabled by the initial deconstruction of population heterogeneity that usually confounds linear modeling of mixed complex data representing multiple regulatory states. These advantages are reflected in the accuracy of the CellOracle simulations we present, and the biological insights they provide.

Our application of CellOracle to the direct reprogramming of MEF to iEPs revealed many new insights into this lineage conversion. First, we discovered two different modes of reprogramming failure. Using CellTag-based lineage tracing, we had previously demonstrated the existence of distinct conversion trajectories: one path leading to successfully reprogrammed cells, and a route to a dead-end state, accompanied by fibroblast gene re-expression (Biddy et al., 2018). From lineage analysis, we found that these trajectories were established at the earliest stages of reprogramming. Here, CellOracle analysis has revealed trajectory-specific GRN configurations, where we observe the high-connectivity of *Zeb1* in cells destined to the dead-end trajectory. Zeb1 is a TF associated with the promotion of epithelial to mesenchymal transition (Liu et al., 2008); thus the induction and expression of this factor may explain the observed persistence of fibroblast marker gene expression, absence of iEP marker expression, and failure to complete mesenchymal to epithelial transition (MET) in dead-end cells.

Cells on the above dead-end trajectory appear to initiate but fail to complete reprogramming. Here, we have uncovered an additional, previously unanticipated dead-end that fails to initiate conversion. Cells on this path express reprogramming transgenes, yet the inferred transgene-target TF connections are significantly weaker, relative to other trajectories. From our lineage analysis, we find cells from a single clone dominate this trajectory. Here, we speculate that the cell giving rise to this clone expresses sufficient levels of transgene, but the target genes required to initiate reprogramming are ‘locked’ within inaccessible heterochromatin, as has been suggested previously (Soufi et al., 2012). Together, these results demonstrate the power of combining lineage analysis, in this instance, via CellTagging (Kong et al., 2020a), with CellOracle network biology analysis to yield mechanistic insight into the different modes of reprogramming failure.

The CellOracle analyses presented here also provide new mechanistic insight into successful reprogramming. Network connectivity scores and cartography analyses suggest the AP-1 subunit *Fos* is a putative reprogramming regulator. Indeed, our simulated perturbations of Fos support its role in the generation and maintenance of iEPs. We confirmed these simulations experimentally, where the addition of *Fos* to the reprogramming cocktail significantly increases the yield of iEPs. Conversely, iEP identity is attenuated upon *Fos* knockout. Further investigation of inferred *Fos* targets implicates a role for Yap1, a Hippo signaling effector, in reprogramming. This observation is supported by our finding that a well-established signature of active Yap1 is enriched as reprogramming progresses, which suggested a role for Yap1, potentially effected via its interactions with Fos/AP-1. Indeed, addition of Fos or Yap1 to the reprogramming cocktail resulted in a significant increase in reprogramming efficiency, where addition of both factors yielded a three-fold increase in iEP colony formation. Intriguingly, in a parallel study, we have found that iEPs resemble post-injury biliary epithelial cells (BECs) (Kong et al., 2020b). Considering that Yap1 plays a central role in liver regeneration (Pepe-Mooney et al., 2019; Yimlamai et al., 2014), these results raise the possibility that iEPs represent a regenerative cell type, explaining their Yap1 activity, self-renewal *in vitro*, and capacity to functionally engraft liver (Sekiya and Suzuki, 2011), and intestine (Guo et al., 2019; Morris et al., 2014). Altogether, these new mechanistic insights have been enabled by CellOracle analysis, placing it as a powerful tool for the dissection of cell identity, aiding improvements in reprogramming efficiency and fidelity.

## Code availability

CellOracle code, documentation, and tutorials are available on GitHub (https://github.com/morris-lab/CellOracle).

## Data availability

All source data, including sequencing reads and single-cell expression matrices, are available from the Gene Expression Omnibus (GEO) under accession codes GSE72859 (Paul et al., 2015) and GSE99915 (Biddy et al., 2018) and GSE145298 for the Perturb-seq data presented in this manuscript.

## Acknowledgments

We thank members of the Morris laboratory for helpful discussions, Jeff Magee, and Todd Druley for critical feedback. This work was funded by National Institute of General Medical Sciences R01 GM126112, and Silicon Valley Community Foundation, Chan Zuckerberg Initiative Grant HCA2-A-1708-02799, both to S.A.M.; S.A.M. is supported by an Allen Distinguished Investigator Award (through the Paul G. Allen Frontiers Group), a Vallee Scholar Award, and a Sloan Research Fellowship; K.K. is supported by a Japan Society for the Promotion of Science Postdoctoral Fellowship.

## Author Contributions

K.K. and S.A.M. conceived the research. K.K. led computational and experimental work, assisted by C.M.H. and supervised by S.A.M. All authors participated in interpretation of data and writing the manuscript.

## Competing Interests

The authors declare no competing interests.

**Correspondence and requests for materials** should be addressed to S.A.M.

## Materials and Methods

### CellOracle overview

CellOracle is an integrative tool for GRN inference and network analysis. It consists of several steps: (1) base GRN construction using scATAC-seq data, (2) context-dependent GRN inference using scRNA-seq data, (3) network analysis, and (4) simulation of cell identity after perturbation. We created the algorithm in Python and also designed it for use in the Jupyter notebook environment. CellOracle code is open source and available on GitHub (https://github.com/morris-lab/CellOracle), along with detailed function descriptions and tutorials.

### (1) Base GRN construction using scATAC-seq data

In the first step, CellOracle constructs a base GRN that contains unweighted, directional edges between a TF and its target gene. For this task, CellOracle utilizes the genomic DNA-sequence of the regulatory region and binding motif sequence of TFs. CellOracle identifies regulatory candidate genes by scanning for TF binding motifs within the regulatory DNA sequences (promoter/enhancers) of open chromatin sites. This process is beneficial as it narrows the scope of possible regulatory candidate genes in advance of model fitting and also helps to define the directionality of regulatory edges in the GRN. However, it is important to note, the base network generated in this step may still contain pseudo- or inactive-connections. This is due to the fact that TF regulatory mechanisms are not only determined by the accessibility of binding motifs, but may also influenced by many context-dependent factors. Thus, scRNA-seq data is used to refine this base network during the model fitting process in the next step.

Base GRN assembly can be divided into two steps: (i) identification of promoter/enhancer regions using scATAC-seq data and (ii) motif scanning of promoter/enhancer DNA sequences.

#### (i) Identification of promoter/enhancer regions using scATAC-seq data

CellOracle uses genomic DNA sequence information to define candidate regulatory interactions. To achieve this, we first need to designate the genomic regions of promoters/enhancers, which we infer from ATAC-seq data. We designed CellOracle for use with scATAC-seq data to identify promoter/enhancers, but in theory, other data should be compatible, such as DNase-seq, ChIP-seq, or Hi-C. We identify promoter/enhancer regions from accessible DNA regions in scATAC-seq data. Thus, scATAC-seq data for a specific tissue/cell-type yields a base GRN representing a sample-specific TF-binding network. In the absence of a sample-specific scATAC-seq dataset, we recommend the use of scATAC-seq data from closely-related tissue/cell-types to support the identification of promoter/enhancer regions. The use of broader scATAC-seq datasets results in a base GRN corresponding to a general TF-binding network, rather than a sample-specific base GRN. Nevertheless, this base GRN network will still be tailored to your specific sample using the scRNA-seq data during the model fitting process, and the final product will consist of context-dependent GRN configurations.

To recognize promoter/enhancer DNA regions within the scATAC-seq data, CellOracle first identifies proximal regulatory DNA elements by locating transcription starts sites (TSS) within the accessible ATAC-seq peaks. This annotation is performed using HOMER (http://homer.ucsd.edu/homer/). Next, the distal regulatory DNA elements are obtained using Cicero, a computational tool that identifies cis-regulatory DNA interactions based on co-accessibility, as dervied from ATAC-seq peak information (Pliner et al., 2018). Cicero identifies a pair of peaks by calculating a co-accessibility score. Then, CellOracle identifies distal cis-regulatory elements that have both a high co-accessibility score (>=0.8) with a DNA element that includes a TSS. These results are saved as a bed file and used in the next step. A database of promoter/enhancer DNA sequences can serve as an alternative if the data is available as a bed file.

#### (ii) Motif Scan of promoter/enhancer DNA sequences

This step scans DNA sequences of promoter/enhancer elements to identify TF binding motifs. CellOracle internally uses gimmemotifs (https://gimmemotifs.readthedocs.io/en/master/), a Python package for TF motif analysis. Here, we use the default gimmemotifs motif database with a false positive rate threshold of 0.02. CellOracle exports a data-table that represents a potential connection between a TF and its target gene, across all TFs and target genes. CellOracle also reports the TF binding DNA region.

### (2) Context-dependent GRN inference using scRNA-seq data

We designed the output of CellOracle GRN inference to be easily interpretable. Because CellOracle leverages genomic sequences and TF binding motif information to infer the base GRN structure and directionality, it does not need to infer causality/directionality of the GRN from gene expression data. This allows CellOracle to adopt a relatively simple machine learning model for GRN inference, a regularized linear machine learning model. CellOracle builds a model that predicts a target gene expression based on the gene expression of regulatory candidate genes:

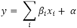

Where *x*_*i*_ is gene expression value of the regulatory candidate gene, *y* is target gene expression, *β*_*i*_ is a coefficient value of the linear model, and *α* is an intercept for this model. Here, we use the list of potential regulatory genes generated in the previous step. In CellOracle, we use the coefficient *β* as a network edge strength between a TF and its target gene. For example, *β* can be high if the target gene expression is highly dependent on TF gene expression. In contrast, *β* can be low if target gene expression does not respond to TF gene expression, even with high expression of both TF and target. Importantly, the gene expression matrix of scRNA-seq data is divided into several clusters in advance by the clustering method such as Louvain clustering or k-means clustering, so that a single data unit for each fitting process should represent a linear relationship, rather than non-linear or mixed regulatory relationships.

CellOracle uses the Bayesian Ridge model or Bagging Ridge model so that we can analyze the reproducibility of the inferred model. In both models, the output is a posterior distribution of coefficient value *β*:

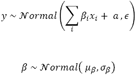

Where μ is the center of the distribution of *β, σ* is the standard deviation of *β*. The user can choose the model method depending on the availability of computational resources and the aim of the analysis; CellOracle’s Bayesian Ridge requires less computational resources, while the Bagging Ridge tends to produce better inference results than Bayesian Ridge. Using the posterior distribution, we can calculate p-values of coefficient *β*; one-sample *t*-tests are applied to *β* to estimate the probability (the center of *β* = 0). The p-value helps identify robust connections while minimizing connections derived from random noise. In addition, we apply regularization to coefficient *β* for two purposes; (i) It is necessary to prevent the coefficient *β* from becoming extremely large due to overfitting, (ii) To identify informative variables via regularization.

In CellOracle, the Bayesian Ridge model uses regularizing prior distribution of *β* as follows:

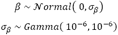

*σ*_*β*_ is selected to represent non-informative prior distributions. With this model, data in the fitting process is used to estimate the optimal regularization strength. In the Bagging Ridge model, arbitrary regularization strength can be manually set.

For the computational implementation of the above machine learning models, we use a Python library, scikit-learn (https://scikit-learn.org/stable/). For Bagging, we use Ridge class in the sklearn.linear_model and BaggingRegressor in the sklearn.ensemble module. The number of iterative calculations in the bagging model can be adjusted depending on the computational resources and amount of time available. For the Bayesian Ridge, we used BayesianRidge class in sklearn.linear_module with the default parameters. Finally, single-cell RNA-seq data requires normalization before the model fitting process. In the default CellOracle pipeline, we recommend performing normalization with the R package, Seurat, using scTransform (https://satijalab.org/seurat/).

### (3) Network analysis

Following GRN inference, we analyze the resulting networks using several graph theory techniques. Before network structure analysis, we filter out weak or insignificant connections. The edge of the GRN is initially filtered based on p-values and absolute values of edge strength. The user can define an arbitrary value for the thresholding according to the data type, data quality, and aim of the analysis. After filtering, CellOracle calculates several network scores: degree centrality, betweenness centrality, and eigenvector centrality. It also assesses network module information and analyzes network cartography. For these processes, CellOracle uses the R packages, igraph (https://igraph.org), linkcomm (https://www.rdocumentation.org/packages/linkcomm/versions/1.0-11), and rnetcarto (https://www.rdocumentation.org/packages/rnetcarto/versions/0.2.4/topics/rnetcarto).

### (4) Simulation of cell identity after regulatory gene perturbation

The central purpose of CellOracle is to understand the mechanism of how a GRN governs cell identity. Toward this goal, we designed CellOracle to leverage inferred GRN configurations to simulate how cell identity changes upon regulatory gene perturbation. To achieve this, CellOracle takes advantage of a linear machine learning model to predict the transition of cell identity after TF perturbation. The simulated gene expression values are converted into a trajectory graph, which represents changes in cell identity, adapting the visualization method previously used by RNA-velocity (La Manno et al., 2018). This process consists of four steps; (i) Data preprocessing, (ii) Signal propagation within the GRN, (iii) Estimation of transition probabilities, (iv) Analysis of simulated transition in cell identity.

#### (i) Data preprocessing

For cell identity simulation, we utilize several functions from Velocyto, a Python package for RNA-velocity analysis (https://velocyto.org). Consequently, CellOracle preprocesses the scRNA-seq data in accordance with Velocyto requirements by first filtering the genes and imputing drop out. Dropout can affect Velocyto’s transition probability calculations, and, thus, KNN imputation must be done before the simulation step. Additionally, CellOracle also constructs a list of candidate TF perturbation targets by selecting 1000 genes with relatively high variability and high gene expression values prior to simulations. Note, we generally avoid simulating the perturbation of a TF if it is not on this list.

#### (ii) Within GRN Signal propagation

This step aims to predict the impact of TF perturbation on cell identity. CellOracle simulates how a cell’s transcriptome changes by utilizing the inferred GRNs. The perturbation effect is then propagated within the GRN to simulate its indirect effects. Here, we take advantage of our inferred GRN because the network edges represent the coefficient values of the linear model. CellOracle uses these GRNs as a model to interpolate/extrapolate target gene expression based on TF gene expression. In this respect, we focus on the differences in gene expression rather than the absolute expression values, so that we can ignore the error or the intercept of the model, which potentially includes unobservable factors within the scRNA-seq data:

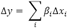

This equation can be converted into matrix multiplication, as follows:

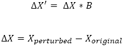

Where *X* is a gene expression matrix (cell x genes), *B* is a matrix of coefficient value *β*. *X*^′^ represents the gene expression matrix in which target gene expression values are updated based on interpolation/extrapolation. We first estimate the effects of perturbation on the first target gene. Then, we calculate the effects of perturbation on the second target genes, based on the calculated difference in the first target gene’s expression. By repeating this calculation for *n* iterations, we can estimate the effects on the n^th^ indirect target gene. During these iterations, the changes caused by the initial perturbation are propagated side-by-side through the connections in the GRNs to simulate indirect global effects. In general, CellOracle performs a small number of iterative calculations (around three cycles) rather than many cycles for two reasons. First, this simulation aims to predict the directionality of the changes in cell identity as opposed to predicting long-term changes in gene expression. A small number of calculations are enough for this task. Second, many iterative calculations may lead to the accumulation of artificial errors, which can distort or exaggerate the results. Of note, CellOracle performs the calculations cluster-wise after splitting the whole gene expression matrix into gene expression submatrices due to the fact that each cluster has a unique GRN configuration. Also, gene expression values are checked between each iterative calculation to confirm whether the extrapolated values exist within a biologically plausible range. If the expression value for a gene becomes negative, this value is adjusted to zero.

#### (iii) Estimation of transition probabilities

From the previous steps, CellOracle produces a simulated gene expression matrix. This gene expression matrix represents future gene expression values following TF perturbation, and the Δ*X* represents the difference of the gene expression. Next, CellOracle aims to project the directionality of the future transition in cell identity onto the dimensional reduction embedding (**Figure 4**). For this task, CellOracle uses a very similar approach to Velocyto. Velocyto visualizes future cell identity based on the RNA-splicing information and calculated vectors from the RNA synthesis and degradation differential equations. CellOracle uses the simulated gene expression matrix, Δ*X*, instead of RNA-velocity vectors. In this process, CellOracle estimates the cell transition probability matrix *P* as follows: *P*_*ij*_ is defined as the probability that cell *i* may adopt a similar cell identity as cell *j* after perturbation. To calculate *P*_*ij*_, CellOracle calculates the Pearson’s correlation coefficient between *d*_*i*_ and *r*_*ij*_:

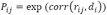

Where *d*_*i*_ is the difference in simulated gene expression (Δ*X*) for cell *i* and *r*_*ij*_ is a difference vector between cell *i* and *j* in the original gene expression matrix. Furthermore, the transition probability P is normalized to fulfill the equation below:

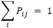

#### (iv) Analysis of simulated cell identity transition

Using the transition probability matrix *P*, CellOracle performs several analyses. First, the transition probability is converted into a transition vector by averaging the local transition probabilities. These calculations and visualizations are done using the same code as Velocyto. Please refer to the Velocyto documentation for more details (http://velocyto.org). CellOracle also performs a Markov simulation to model cell identity transitions. For this simulation, the original cell identity defines the space of possible cell identities. CellOracle uses an original, unperturbed cell identity to define the initial state. The next state is simulated based on the current state and transition probability matrix *P*. The transition process is repeated iteratively, allowing CellOracle to estimate how the identity of a cell changes following perturbation. These results are visualized as a Sankey diagram.

### 10x alignment, digital gene expression matrix generation

The Cell Ranger v2.1.0 pipeline (https://support.10xgenomics.com/single-cell-gene-expression/software/downloads/latest) was used to process data generated using the 10x Chromium platform. Cell Ranger processes, filters, and aligns reads generated with the Chromium single-cell RNA sequencing platform. This pipeline was used in conjunction with a custom reference genome, created by concatenating the sequences corresponding to the *Hnf4α-t2a-Foxa1* transgene as a new chromosome to the mm10 genome. The unique UTRs in the *Hnf4α-t2a-Foxa1* transgene construct allowed us to monitor transgene expression. To create Cell Ranger compatible reference genomes, the references were rebuilt according to instructions from 10x (https://support.10xgenomics.com/single-cell-gene-expression/software/pipelines/latest/advanced/references). To achieve this, we first created a custom gene transfer format (GTF) file, containing our transgenes, followed by indexing of the FASTA and GTF files, using Cell Ranger ‘mkgtf’ and ‘mkref’ functions. Following this step, the default Cell Ranger pipeline was implemented, then the filtered output data used for downstream analyses.

## Experimental Methods

### Mice and derivation of mouse embryonic fibroblasts

Mouse Embryonic Fibroblasts were derived from E13.5 C57BL/6J embryos. (The Jackson laboratory: 000664). Heads and visceral organs were removed from E13.5 embryos. The remaining tissue was minced with a razor blade and then dissociated in a mixture of 0.05% Trypsin and 0.25% Collagenase IV (Life Technologies) at 37°C for 15 minutes. After passing the cell slurry through a 70μM filter to remove debris, cells were washed and then plated on 0.1% gelatin-coated plates, in DMEM supplemented with 10% FBS (Sigma-Aldrich), 2mM L-glutamine, and 50mM β-mercaptoethanol (Life Technologies). All animal procedures were based on animal care guidelines approved by the Institutional Animal Care and Use Committee.

### Retrovirus Production

Retroviral particles were produced by transfecting 293T-17 cells (ATCC: CRL-11268) with the pGCDN-Sam construct containing Hnf4α-t2a-Foxa1/Fos/Zfp57/Yap1, along with packaging construct pCL-Eco (Imgenex). Virus was harvested 48hr and 72hr after transfection and applied to cells immediately following filtering through a low-protein binding 0.45μM filter.

### Generation and collection of iEPs

Mouse embryonic fibroblasts (younger than passage 6) were converted to iEPs as in (Biddy et al., 2018), modified from (Sekiya and Suzuki, 2011). Briefly, we transduced cells every 12hr for 3 days, with fresh Hnf4α-t2a-Foxa1 retrovirus, in the presence of 4mg/ml Protamine Sulfate (Sigma-Aldrich), followed by culture on 0.1% gelatin-treated plates for 1 week in hepato-medium (DMEM:F-12, supplemented with 10% FBS, 1 mg/ml insulin (Sigma-Aldrich), dexamethasone (Sigma-Aldrich), 10mM nicotinamide (Sigma-Aldrich), 2mM L-glutamine, 50mM β-mercaptoethanol (Life Technologies), and penicillin/streptomycin, containing 20 ng/ml hepatocyte growth factor (Sigma-Aldrich), and 20 ng/ml epidermal growth factor (Sigma-Aldrich). After the seven days of culture, the cells were transferred onto plates coated with 5μg/cm^2^ Type I rat collagen (Gibco, A1048301). For single-cell processing, 30,000 reprogrammed, expanded iEPs were collected and fixed in methanol, as previously described in (Alles et al., 2017). Briefly, cells were collected and washed in Phosphate Buffered Saline (PBS), followed by resuspension in ice-cold 80% Methanol in PBS, with gentle vortexing. These cells were stored at -80°C for up to three months, and processed on the 10x platform (below).

### Fos/Zfp57/Yap1 reprogramming and colony formation assays

Mouse *Fos, Zfp57*, and *Yap1* were cloned from iEPs into the retroviral vector, pGCDNSam (Sekiya and Suzuki, 2011), and retrovirus produced as above. For comparative reprogramming experiments, mouse embryonic fibroblasts (2×10^5^/well of a 6-well plate) were serially transduced over 72hr (as above). In control experiments, virus produced from an empty vector control expressing only GFP was added to the Hnf4α-Foxa1 reprogramming cocktail. In Fos, Zfp57, or Yap1 experiments, virus produced from the Fos/Zfp57/Yap1-IRES-GFP constructs was added to Hnf4α and Foxa1. Fos/Zfp57/Yap1 overexpression was confirmed by harvesting RNA from Hnf4α-Foxa1 and Hnf4α-Foxa1-Fos/Zfp57/Yap1-transduced cells (RNeasy kit, Qiagen). Following cDNA synthesis (Maxima cDNA synthesis kit, Life Tech), qPCR was performed to quantify *Fos/Zfp57/Yap1* overexpression (TaqMan Probes: Gapdh Mm99999915_g1; *Cdh1* Mm01247357_m1; *Apoa1* Mm00437569_m1; *Fos* Mm00487425_m1; *Yap1* Mm01143263_m1; *Zfp57* Mm00456405_m1, TaqMan qPCR Mastermix, Applied Biosystems). Cells underwent reprogramming for two weeks and were processed for colony formation assays: cells were fixed on the plate with 4% PFA, permeabilized in 0.1% Triton-X100 then blocked with Mouse on Mouse Elite Peroxidase Kit (Vector PK-2200). Primary antibody, mouse anti-E-Cadherin (1:100, BD Biosciences) was applied for 30 min before washing and processing with the VECTOR VIP Peroxidase Substrate Kit (Vector SK-4600). Colonies were visualized on a flatbed scanner, adding heavy cream to each well in order to increase image contrast. Colonies were counted, using our automated colony counting tool: https://github.com/morris-lab/Colony-counter.

### Perturb-seq

We performed Perturb-seq as previously described (Adamson et al., 2016). The protocol was modified, as outlined below, to apply the strategy to our experimental system:

#### (1) Vector backbone and gene barcode pool construction

For Perturb-seq experiments, we used a lentivirus vector to express guide RNAs and gene barcodes (GBC). The lentivirus vector backbone contains an antiparallel cassette containing a guide RNA and GBC. In the original perturb-seq paper, the authors used pPS and pBA439 to construct the guide RNA-GBC vector pool. Here, we modified pPS and pBA439 to generate the pPS2 vector, in which the Puromycin-t2a-BFP gene was replaced by the Blasticidin-t2a-BFP gene. We constructed the guide RNA-GBC vector using a multi-step cloning strategy: First, we synthesized dsDNA, via PCR, for a random GBC pool. We purified the PCR product with AMPure XP SPRI beads. We then inserted the purified GBC pool into the pPS2 vector at the EcoRI site in the 3’ UTR of the Blasticidn-t2a-BFP gene. We used the product of Gibson assembly for transformation into DH5α competent cells (NEB: C2987H). Transformed cells were cultured directly in LB liquid. We extracted plasmid DNA to yield the pPS2-GBC pool.

#### (2) Guide RNA cloning

We designed guide RNAs using https://zlab.bio/guide-design-resources. We synthesized oligo DNA for each guide RNA. Oligo DNA pairs were annealed and inserted into the pPS2-GBC vector, following BsmB1 digestion. After isolation and growth of single colonies, plasmid DNA was extracted and sanger DNA-sequenced; sequences of the guide RNA inserted site and GBC site were used to construct a gRNA/GBC reference table:

Fos_sg0 CAGCCGACTGAACGCGTTATTC

Fos_sg1 CATATATCAAAGATGAACATTG

Fos_sg2 TCAAGGCTGTAATTTCTTGGGC

empty0 TTGATGAACTGCGCTAGCGAGG

empty1 AAGAGCGGCTCGCAAGGGAAAA

empty2 AGTAGGATACGTGGAGTTAATA

#### (3) Lentivirus guide RNA pool generation

An equal amount of DNA for each pPS2-guide RNA vector was mixed together to generate the plasmid pool. Three control vectors were also mixed with this plasmid vector pool; the weight ratio of each pPS2-guide vector to each control vector was 1:4. We used this mixed DNA pool for lentivirus production. Lentiviral particles were produced by transfecting 293T-17 cells (ATT: CRL-11268) with the pPS-guide RNA-GBC constructs, along with the packaging plasmid, psPAX2 (https://www.addgene.org/12260/), and pMD2.G (https://www.addgene.org/12259/).

#### (4) Cell culture for Perturb-seq

We transduced reprogrammed iEP cells with retrovirus carrying Cas9 (MSCV-Cas9-Puro). The cells were treated with Puromycin (4 μg/ml) for four days to eliminate non-transduced cells. iEP-Cas9 cells were transduced with the lentivirus guide RNA pool for 24 hours. The concentration of lentivirus was pre-determined to target 10∼20% transduction efficiency. After four days of cell culturing, we sorted BFP positive cells to purify transduced cells. Cells were cultured for a further 72 hours and fixed with methanol as previously described (Alles et al., 2017).

#### (5) GBC amplification and sequencing

Following library preparation on the 10x chromium platform (below), we PCR amplified the GBC. The amplification was performed largely according the original perturb-seq paper (Adamson et al., 2016), but we modified the PCR primer sequence for the Chromium single cell library v2 kit:

P7_ind_R2_BFP_primer: CAAGCAGAAGACGGCATACGAGATTCGCCTTAGTGACTGGAGTTCAGACGTGTGCTCTTCCGATCTTAGCAAACTGGGGCACAAGC

P5_partial_primer: AATGATACGGCGACCACCGA

GBG_Amp_F: GCTGATCAGCGGGTTTAAACGGGCCCTCTAGG

GBG_Amp_R: CGCGTCGTGACTGGGAAAACCCTGGCGAATTG

GBC_Oligo: TTAAACGGGCCCTCTAGGNNNNNNNNNNNNNNNNNNNNNNCAATTCGCCAGGGTTTTCCC

Following amplification, we purified the PCR product with AMPure XP SPRI beads. The purified sample was sequenced on the Illumina Mi-seq platform.

#### (6) Alignment of cell barcode/GBC

For preprocessing of Perturb-seq metadata, we used MIMOSCA, a computational pipeline for the analysis of perturb-seq data (https://github.com/asncd/MIMOSCA). First, the reference table for the cell barcode/GBC pair was generated from Fastq files. The data table was converted into the guide RNA/cell barcode table using the guide RNA-GBC reference table. This metadata was integrated into the scRNA-seq data. The guide metadata was processed with an EM-like algorithm in MIMOSCA to filter out unperturbed cells computationally as previously described (Adamson et al., 2016).

### 10x procedure

For single-cell library preparation on the 10x Genomics platform, we used: the Chromium Single Cell 3′ Library & Gel Bead Kit v2 (PN-120237), Chromium Single Cell 3′ Chip kit v2 (PN-120236) and Chromium i7 Multiplex Kit (PN-120262), according to the manufacturer’s instructions in the Chromium Single Cell 3′ Reagents Kits V2 User Guide. Just prior to cell capture, methanol-fixed cells were placed on ice, then spun at 3000rpm for 5 minutes at 4°C, followed by resuspension and rehydration in PBS, according to (Alles et al., 2017). 17,000 cells were loaded per lane of the chip, aiming to capture 10,000 single-cell transcriptomes. Resulting cDNA libraries were quantified on an Agilent Tapestation and sequenced on an Illumina HiSeq 2500.

## Supplementary Figure Legends

**Supplementary Figure 1 (Related to Figure 2).**
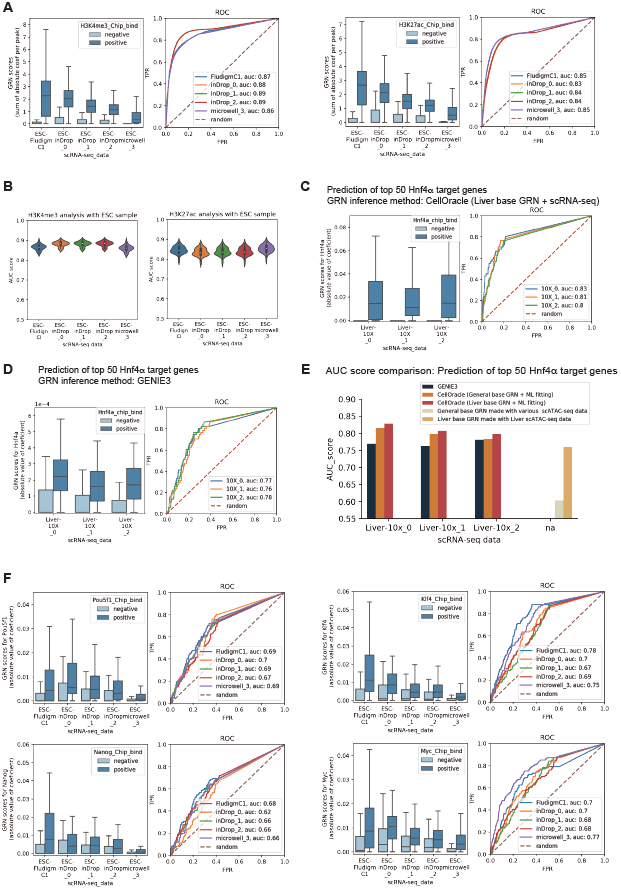
Benchmarking, and validation of inferred GRN configurations. **(A)** Comparison of inferred promoter/enhancer element activity between positive and negative ChIP-seq peaks for active histone marks in embryonic stem cells (ESCs); Upper panels: H3K4me3; Lower panels: H3K27ac. Box plots, dark blue: positive peaks; light blue: negative peaks. The y-axis of the box plot is the inferred GRN score for each DNA peak. The score is defined as the sum of the absolute value of the GRN connection at the peak. Outliers, determined by the interquartile outlier rule, are not shown). ROC curves show predictive scores when we predict H3K4me3 peaks and H3K27ac peaks based on the GRN score. **(B)** Comparison of AUC scores from **(A)** across datasets and platforms. **(C)** CellOracle benchmarking against GENIE3: Comparison between inferred Hnf4α target genes and Hnf4α ChIP-seq experimental data for the liver. For GRN inference here, CellOracle used a Liver scATAC-seq dataset from the mouse scATAC-seq atlas dataset (Cusanovich et al., 2018) to generate a base GRN. **(D)** GENIE3 was applied to the three 10x Liver scRNA-seq data to infer Liver GRNs. **(E)** Comparison of CellOracle and GENIE3 AUC scores. (**F**) Comparison between inferred regulatory connections and ChIP-seq experimental data for several TFs in ESCs: Pou5f1/Oct4, Klf4, Nanog, Myc, and Sox2. CellOracle’s GRN configuration inference algorithm was applied to scRNA-seq data to predict active target genes for each reprogramming TF. The y-axis of the box plot shows the inferred score; the absolute mean coefficient values in the bagging ridge model. The top 100 target genes, based on the rank of MACS2 binding scores, are shown in dark-blue. Non-target genes are shown in light blue. Outliers, which were determined by the interquartile outlier rule, are not shown in the box plot. The ROC curve shows prediction scores when we predict the top 100 target genes for each TF based on the strength of the GRN score. The scRNA-seq datasets used here include several different platforms: Fludigm-C1, inDrop, and Microwell-seq (**Table S1**).

**Supplementary Figure 2 (Related to Figure 3).**
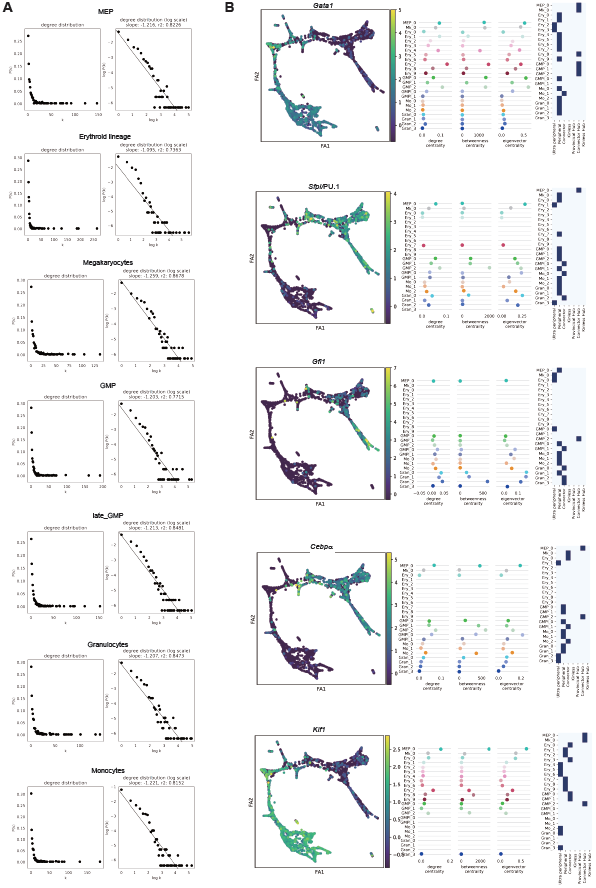
(A) Degree distribution of HSC GRN configurations. CellOracle inferred cell-type-specific GRN configurations for the Paul et al., 2015 dataset. We calculated degree distribution for each GRN configuration after pruning weak network edges, based on the p-value and strength; Edges with a p-value more than 0.001 were excluded, and then the top 2000 edges with high absolute mean coefficient values were selected. We counted the network degree (k), representing the number of network edges for each gene. P(k) is the frequency of network degree k. The relationship between k and P(k) were visualized in a scatter plot. We also visualized this relationship after log-transformation to test whether the network is a ‘scale-free network.’ The degree distribution of a scale-free network follows a power law; there is a linear relationship between log(k) and log(P(k)). These plots demonstrate that these are indeed scale-free networks. **(B) Network score for selected genes within HSC GRN configurations.** We calculated network scores and gene cartography roles for TFs that are known to play cell-type-specific roles in hematopoiesis. The first column shows the projection of gene expression onto the force-directed graph. The second column shows network scores of the gene of interest for each GRN configuration. The third column shows the cartography role for each cluster-specific GRN configuration. If the TF has no network edge after filtering, the network score and cartography data point is empty.

**Supplementary Figure 3 (Related to Figure 3).**
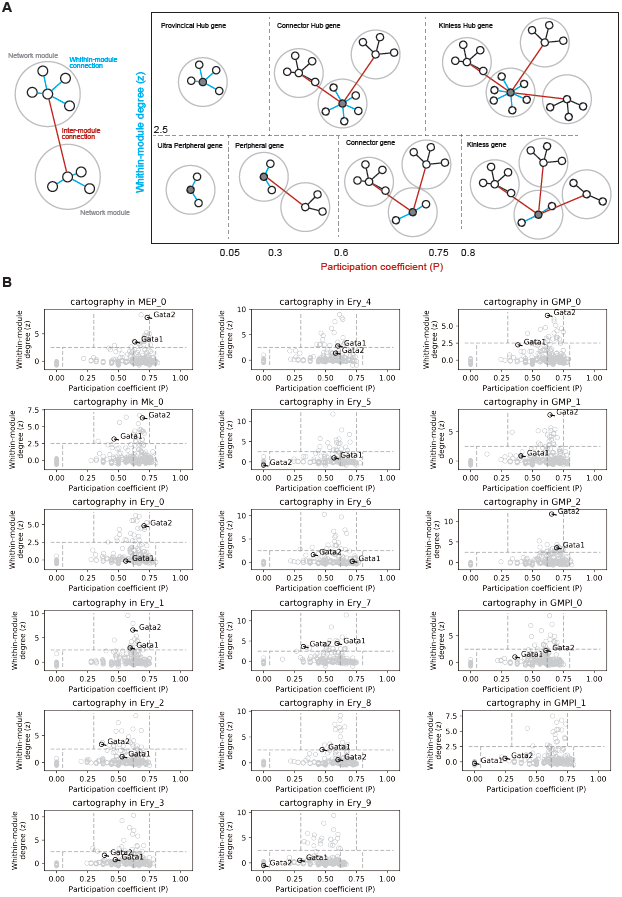
Cartography analysis with HSC GRN configurations. **(A)** Illustration of the cartography analysis method. The cartography method classifies genes into seven groups according to two network scores: within-module degree and participation coefficient. We calculated these values using the CellOracle inferred GRN configurations (see methods). **(B)** The results of cartography analysis with hematopoietic GRN configurations. We calculated the within-module degree and participation coefficient for each cluster-specific GRN configuration.

**Supplementary Figure 4 (Related to Figure 4).**
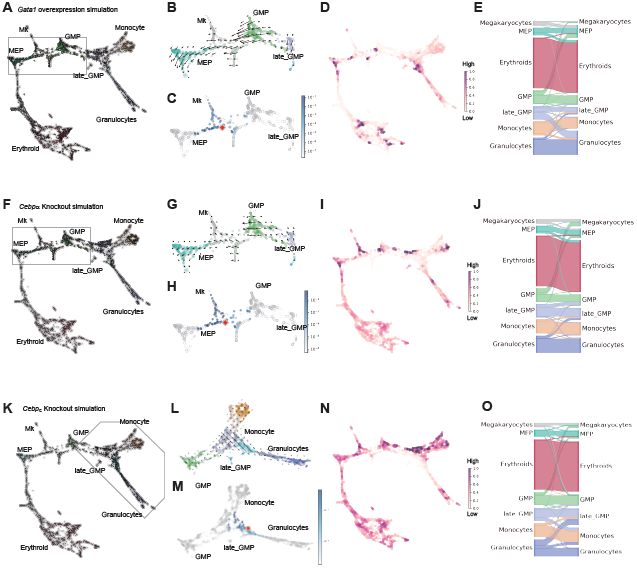
Additional simulation with HSC GRN configurations. We simulated cell state transitions resulting from the perturbation of specific TFs. We performed simulations for *Gata1* overexpression (**A-E**), *Cebpα* knock-out (**F-J**), and *Cebpε* knock-out (**K-O**). For the simulation of *Gata1* overexpression, we set the *Gata1* expression value at 0.930, which is the maximum value of *Gata1* expression in the imputed gene expression matrix. **(A/F/K)** The plot shows the CellOracle estimation of cell type transition after *Gata1* overexpression. **(B, C/G, H/L, M)** The cell trajectory graph around MEP, GMP, late_GMP clusters, is magnified to show the cell transition in these clusters. This area is highlighted by a light gray line in panel A. **(B/G/L)** The graph shows the local average of the transition vector on the grid. **(C/H/M)** Cell transition probability score of one specific cell, shown as a red diamond (representing a midpoint between MEP and GMP identity). CellOracle simulated cell transition by a Markov simulation method, using the cell transition probability (number of simulation steps = 100). **(D/I/N)** Cell density after the simulation, projected onto the force-directed graph. **(E/J/O)** Sankey diagram providing a summary of the cell transition simulation, with cells grouped by cluster. For the simulation of *Cebpα* knock-out, we set the *Cebpα* expression at 0. The other simulation parameters were the same as above. For the simulation of *Cebpε* knock-out, we set the *Cebpε* expression at 0. The other simulation parameters were the same as above.

**Supplementary Figure 5 (Related to Figure 5).**
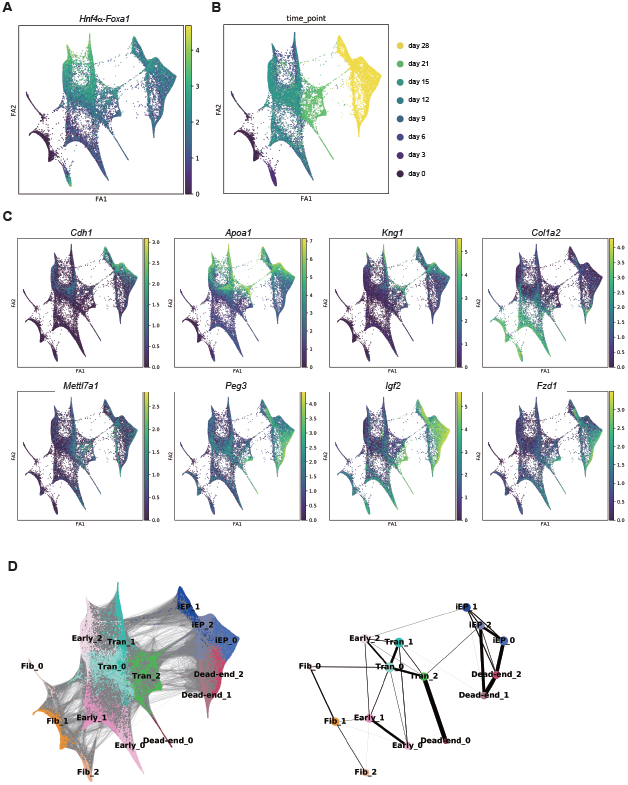
Single-cell analysis of fibroblast to iEP reprogramming. Dimensionality reduction plot of iEP reprogramming scRNA-seq data made with force-directed graph drawing. Cells classify into 15 groups by the Louvain clustering method. **(A)** Projection of *Hnf4α-t2a-Foxa1* (*Hnf4α-Foxa1)* transgene expression levels onto the force-directed graph **(B)** Projection of reprogramming time point information onto the force-directed graph. There are 8 time points; day 0, 3, 6, 9, 12, 15, 21, and 28. **(C)** Projection of gene marker expression for the phases/cell types arising during iEP reprogramming. Reprogrammed iEP cell cluster marker genes: *Cdh1, Apoa1*, and *Kng1*. Fibroblast marker gene: *Col1a2*. Transition marker gene: *Mettl7a1*. Dead-end marker genes: *Peg3, Igf2*, and *Fzd1.* **(D)** Partition-based graph abstraction (PAGA) (Wolf et al., 2019) analysis of 27,663 reprogramming cells from reveals 15 major clusters, reproducing the known conversion trajectories. Cell clusters were annotated manually with marker gene expression and grouped into five cell types; Fibroblasts, Early_Transition, Transition, Dead-end, and Reprogrammed iEP.

**Supplementary Figure 6 (Related to Figure 5).**
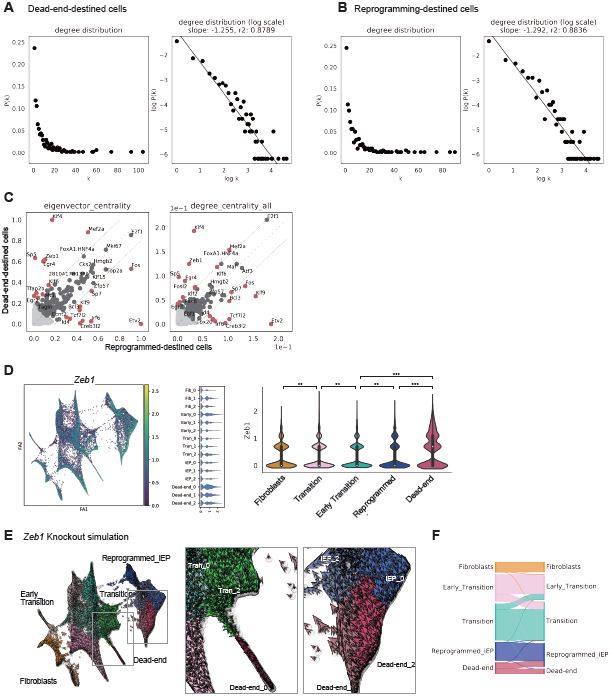
CellOracle network analysis of cells destined to reprogrammed or dead-end states. **(A, B)** We identified early-stage cells (days 6-15) on the reprogrammed (iEP_0, iEP_1, and iEP_2 clusters), and dead-end (Dead-end_1, and Dead-end_2 clusters) trajectories. For this, we aggregated cells across days 6-15 to increase the power of our analysis, resulting in n = 51 dead-end cells, and n = 42 reprogrammed cells. For these two groups, we used CellOracle to infer GRN configurations, first assessing the quality of the inferred networks. We calculated degree distribution for each GRN configuration after pruning weak network edges, based on the p-value and strength, as above (**Figure S2A**). We counted the network degree (k), representing the number of network edges for each gene. P(k) is the frequency of network degree k, visualized in scatter plots (left panels, **A, B**). We also visualized the relationship between k and P(k) after log-transformation shows that these are scale-free networks (right panels, **A, B**), demonstrating successful network inference from these relatively small cell populations. **(C)** Comparison of eigenvector and degree centrality score between the GRN configurations of dead-end and reprogramming-destined cells. **(D)** *Zeb1* expression projected onto the force-directed graph (left panel). Violin plot showing *Zeb1* expression across all clusters (middle panel). Violin plot showing *Zeb1* expression, grouped by cluster type (** = *P* <0.01, *** = *P* <0.01, permutation test, one-sided). **(E)** *Zeb1* knockout simulation. The plot (left panel) shows the CellOracle estimation of cell type transition after *Zeb1* knockout, with *Zeb1* expression set to 0. Right panels: magnified areas outlined in left panels. **(F)** Sankey diagram showing the number of cell transitions between clusters.

**Supplementary Figure 7 (Related to Figure 6):**
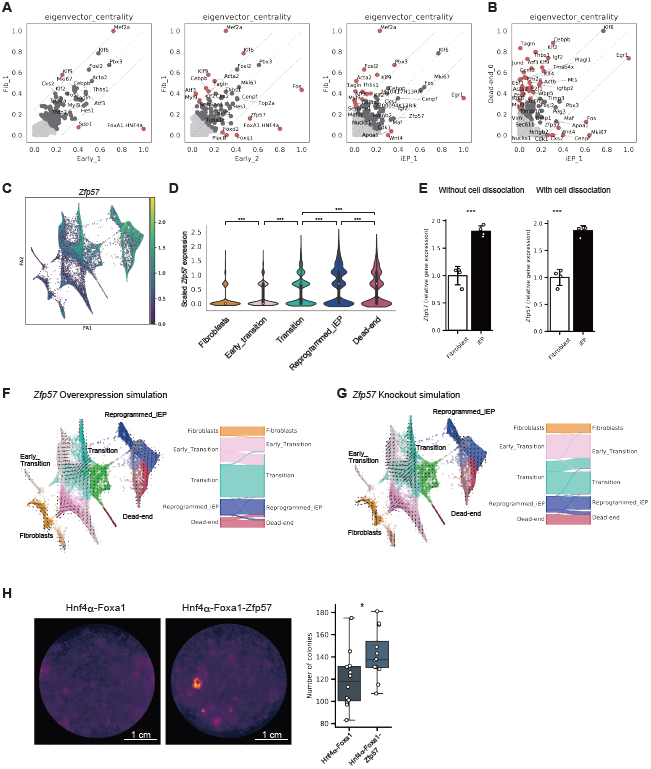
CellOracle analysis of the role of Zfp57 in fibroblast to iEP reprogramming. **(A, B)** Scatter plots showing a comparison of eigenvector centrality scores between specific clusters. **(A)** Comparison of eigenvector centrality score between the Fib_1 cluster GRN configuration and the GRN configurations of other clusters in relatively early stages of reprogramming. **(B)** Comparison of eigenvector centrality score between iEP_1 and Dead-end_0 cluster GRN configurations. **(C)** *Zfp57* expression projected onto the force-directed graph of fibroblast to iEP reprogramming. **(D)** Violin plot of *Zfp57* expression across reprogramming stages. **(E)** qPCR of *Zfp57* expression in fibroblasts and iEPs, with and without cell dissociation prior to the assay. **(F)** *Zfp57* gene overexpression simulation with reprogramming GRN configurations. The left panel is the projection of simulated cell transitions onto the force-directed graph. The Sankey diagram summarizes the simulation of cell transitions between cell clusters. For the simulation of *Zfp57* overexpression, we set the *Zfp57* expression value at 0.9649, which is the maximum value of *Zfp57* expression in the imputed gene expression matrix. **(G)** *Zfp57* gene knockout simulation. **(H)** Colony formation assay with addition of *Zfp57* to the Hnf4α-Foxa1 reprogramming cocktail. Left panel: E-cadherin immunohistochemistry. Right panel: box plot colony numbers (n = 6 technical replicates, 2 independent biological replicates; * = *P* < 0.05, *t-*test, one-sided).

**Supplementary Figure 8 (Related to Figure 6).**
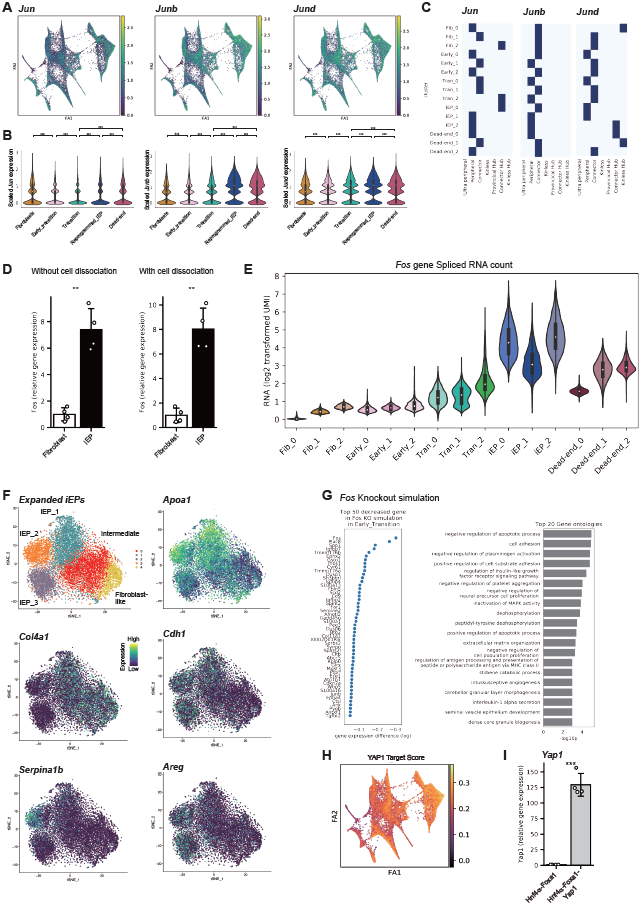
CellOracle analysis of the role of Fos and Yap1 in fibroblast to iEP reprogramming. **(A-C)** Expression and network cartography of Jun family members, *Jun, Junb*, and *Jund*. **(D)** qPCR of *Fos* expression in fibroblasts and iEPs, with and without cell dissociation prior to the assay, ** = *P* < 0.01, *t-*test, one-sided. **(E)** Analysis of *Fos* mRNA splicing state in the scRNA-seq data of iEP reprogramming to investigate the *Fos* mRNA maturation state: Violin plot for spliced *Fos* mRNA counts. **(F)** *t*-SNE plots of 9,914 expanded iEPs, cultured long-term, revealing fibroblast-like, intermediate, and three iEP subpopulations. Expression levels of *Apoa1* (marking typical iEPs), *Col4a1* (fibroblast-like cells), *Cdh1, Serpina1b* (hepatic-like iEPs), and *Areg* (intestine-like iEPs) projected onto the *t*-SNE plot. Top 50 decreased genes in *Fos* knockout simulation in the early reprogramming transition (left) and GO analysis based on these genes (right). **(H)** Projection of YAP1 target gene scores onto the force directed graph of reprogramming. **(I)** qPCR assay for *Yap1* expression following addition of Yap1 and Fos to the Hnf4α-Foxa1 reprogramming cocktail (n = 4 biological replicates; *** = *P* < 0.001, ** = *P* < 0.01, *t-*test, one-sided), confirming Yap1 overexpression.

**Supplementary Table 1.** Details of the publicly available scRNA-seq and ChIP-seq datasets used in this study.

**Supplementary Table 2.** Top 50 CellOracle-inferred *Fos* targets, across all reprogramming clusters. Confirmed YAP1 targets are highlighted in red.

